# MKPV (aka MuCPV) and related chapparvoviruses are nephro-tropic and encode novel accessory proteins p15 and NS2

**DOI:** 10.1101/732537

**Authors:** Christopher J. Jolly, Quintin Lee, Matthew P. Padula, Natalia Pinello, Simon H. Williams, Matthew B. O’Rourke, Marcilio Jorge Fumagalli, Joseph D. Orkin, Babak Shaban, Ori Brenner, Wolfgang Weninger, William Marciel de Souza, Amanda D. Melin, Justin J.-L. Wong, Marcus J. Crim, Sébastien Monette, Ben Roediger

## Abstract

Mouse kidney parvovirus (MKPV) is a member of the provisional *Chapparvovirus* genus that causes renal disease in immune-compromised mice, with a disease course reminiscent of polyomavirus-associated nephropathy in immune-suppressed kidney transplant patients. Here we map four MKPV transcripts, created by alternative splicing, to a common transcription initiation region, and use mass spectrometry to identify “p10” and “p15” as novel chapparvovirus accessory proteins produced in MKPV-infected kidneys. p15 and a splicing-dependent putative accessory protein NS2 are conserved in all near-complete tetrapod chapparvovirus genomes currently available (from mammals, birds and a reptile). In contrast, p10 may be encoded only by viruses with >60% amino acid identity to MKPV. We show that MKPV is kidney-tropic and that the bat chapparvovirus DrPV-1 and a non-human primate chapparvovirus, CKPV, are also found in the kidneys of their hosts. We propose, therefore, that chapparvoviruses with >60% VP1 amino acid identity to MKPV be classified into a genus dubbed *Nephroparvovirus*, which is consistent with nomenclature for the genus *Erythroparvovirus*.

## Introduction

Parvoviruses are small, non-enveloped, polyhedral, single-strand DNA viruses with genomes 4– 6kb in length which bear short (120–600 base) inverted terminal repeats (ITRs) forming hairpin telomeres. All parvoviruses comprise 2 major genes encoding a non-structural replication protein NS1 (gene *rep*) and a capsid protein VP1 (gene *cap*). Alternative splicing or alternative translation initiation sites can allow the production of truncated forms of VP1; all sharing the same C-terminal region (1, 2). Open reading frames (ORFs) overlapping the major NS1 or VP1 reading frames encode smaller genus-specific accessory proteins. Genetic simplicity combined with a single-stranded DNA genome dictates that parvoviruses can only replicate when the host cell itself replicates. Many members of the *Dependoparvovirus* genus (e.g. adeno-associated virus, AAV) can furthermore only replicate if a helper virus is also present (1, 3), but this is not a universal feature of the *Dependoparvovirus* genus – close avian relatives of AAV that cause Derzsy’s disease in geese and Muscovy ducks replicate autonomously (4).

Vertical transmission of parvoviruses across the placenta can kill developing embryos or newborns in domesticated species such as dogs and pigs (5, 6), but many parvoviruses are highly adapted to infecting specific cell types. For instance, *Erythroparvovirus* B19 infects red blood cell precursors in humans, potentially inducing anaemia (7), and even though AAV2 can transduce many tissues, it mostly targets the liver if intravenously injected and is naturally liver-adapted (8). Horizontal transmission of the newly-identified mouse kidney parvovirus (MKPV) induces adult renal failure in severely immune-deficient laboratory mice, without obvious pathology in other tissues (9). Co-incidentally, a virus very similar to MKPV was identified in mice living wild in New York City (NYC) and dubbed murine chapparvovirus (MuCPV), but the state of kidney disease was not assessed in that study (10).

MKPV and MuCPV are only distantly related to other known murine parvoviruses and are members of the provisional genus *Chapparvovirinae*; so-called because the earliest examples, discovered by metagenomic analyses, were found in <u>ch</u>iropteran, <u>a</u>vian and <u>p</u>orcine hosts (i.e. bats, birds and pigs) (11-13). Recently, additional chapparvovirus sequences were discovered by screening of draft genome assemblies; presumed to reflect parvoviral infection of the source animal rather than viral genome integration (14). The growing list of host species now includes marsupials, fish and invertebrates, with chapparvoviral genomes incorporated into some invertebrate genomes (15-17). Thus, chapparvoviruses are an ancient lineage within the family *Parvoviridae*, clustering separately from members of the two currently established parvoviral subfamilies *Parvovirinae* and *Densovirinae* (16). Curiously, fish and arthropod chapparvoviruses are more related to each other than to tetrapod chapparvoviruses (16). In all, at least forty-seven chapparvovirus species have been identified so far; in the tetrapods spread across bats, pigs, rodents, dogs, non-human primates, marsupials, birds and reptiles.

MKPV is currently the only chapparvovirus for which the inverted terminal repeats (ITR) are published, and the only one proved to be viable, infective and pathogenic, to date. Here, we extend our characterisation of MKPV and related viruses. Specifically, we assess the global distribution of MKPV and report co-occurrence of MKPV with mouse kidney disease at additional sites, analyse MKPV tropism, map the major MKPV transcripts, describe a closely-related full-length chapparvovirus from a primate kidney, and identify accessory proteins in MKPV that are conserved in tetrapod-hosted *Chapparvovirinae* genomes. Conservation of genome structure, coding potential and kidney association suggest that many mammal-infecting chapparvovirus species may be adapted to a nephron niche.

## Results

### MKPV and MuCPV are the same virus species

The *Rep* plus *Cap* sequence of “murine chapparvovirus” (MuCPV), lacking ITRs, was originally assembled from the faecal virome of house mice living wild in New York City (NYC; accession MF175078) (10). Independently, a full-length 4,442 nt sequence of “mouse kidney parvovirus” (MKPV), including ITRs, was assembled from the kidney viromes of two renal disease-affected immune-deficient *Rag1*^*-/-*^ mice in the colony of the Centenary Institute, Sydney, Australia (CI; accession MH670587), and a 3.5 kb fragment of MKPV encompassing NS1 and VP1 was then amplified by PCR from the kidneys of two immune-deficient mice necropsied at Memorial Sloan Kettering Cancer Center, NYC (MSKCC; accession MH670588) (9). BLAST revealed that the MuCPV and MKPV genomes are 98% identical to one another; thus, they belong to the same species according to ICTV guidelines. Using primers 889, 890, 891 and 893 based on the CI-MKPV strain (Fig 1A and Table S1), the 5’- and 3’-sequences of NYC-MuCPV and MSKCC-MKPV were extended outwards to include the innermost repeat and hairpin region of each ITR (see NCBI accessions MF175078.2 and MH670588). This confirmed that MuCPV and MKPV share identical heterotelomeric ITRs (Fig 1A).

**Figure 1.**
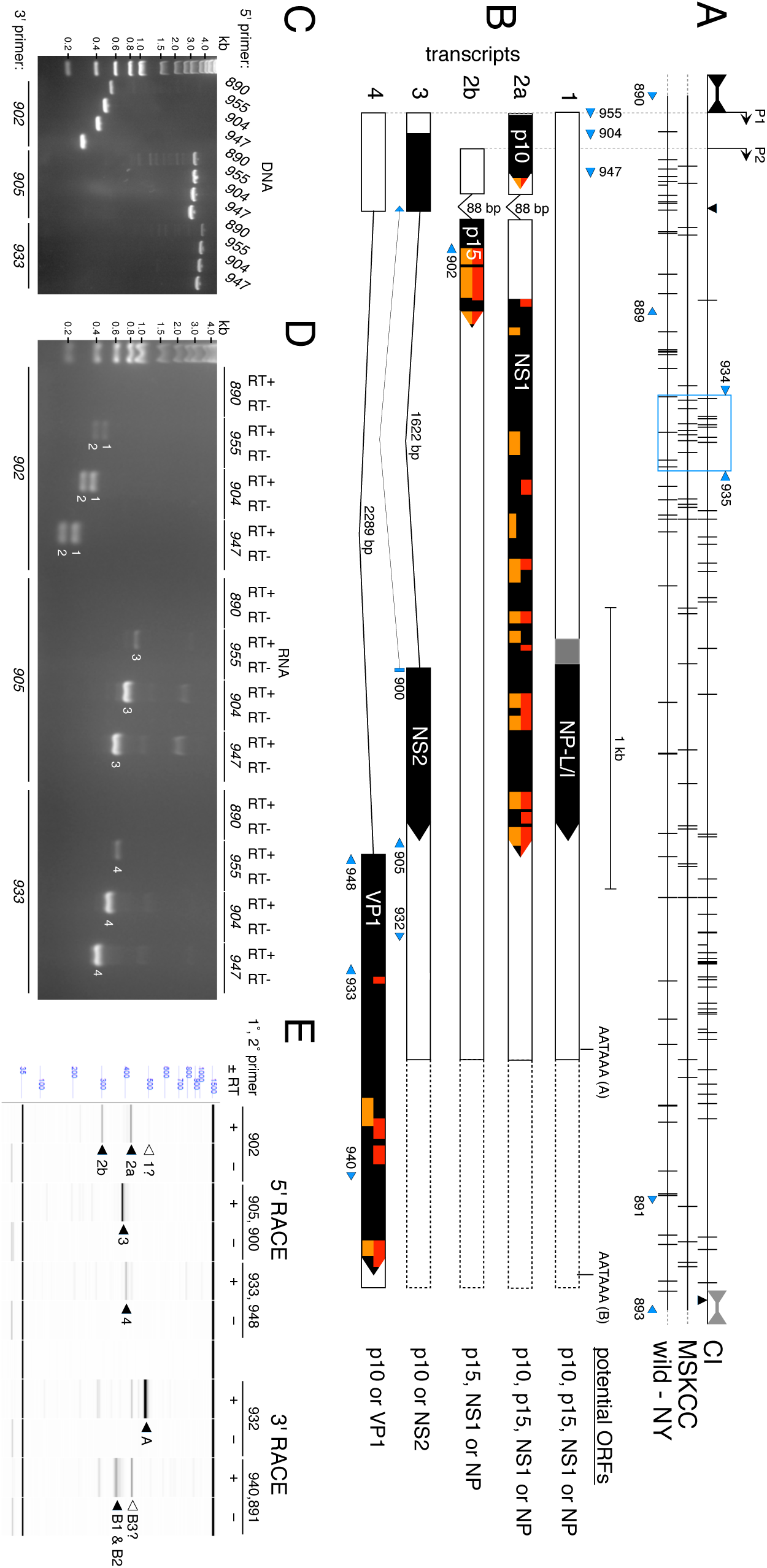
Structure of the MKPV genome. **(A-B)** Maps of the MKPV/MuCPV strains from Centenary Institute (CI, accession MH670587.1), Memorial Sloan Kettering Cancer Center (MSKCC, accession MH670588.2) and New York City basements (wild-NY, MF175078.2). “Bowties” indicate ITRs. **(A)** SNVs between the CI, MSKCC and wild-NY accessions. Vertical lines - differences between accessions. Half height vertical lines - polymorphisms within an accession. Down-pointing triangle – 2 bp insertion in the CI strain. Up-pointing triangle – 1 bp insertion in a CI sub-strain. Dashed lines - missing extremities in MSKCC and wild-NY accessions, which consist of the exterior inverted repeats in the full-length CI sequence. **(B-C)** Alternative splicing allows production of the polypetides p15, NS1, NS2-P and VP1. Red or orange indicate peptides present in LC-MS/MS datasets PXD010540 (9) or PXD014938 (this paper), respectively. p15, p10 and NP-L/I could theoretically be produced from multiple transcripts. **(C-D)** Detection of spliced transcripts by RT-PCR, using primers mapped in A-B. Input templates were MKPV-infected **(C)** kidney DNA or **(D)** DNAse/ExoI-treated RNA, converted (RT+) or mock-converted (RT-) to cDNA. RT-PCR products corresponding to transcripts 1 to 4 are indicated by white numbers. **(E)** Mapping of transcription start and stop sites by RACE. See Fig S1 for RACE details. Major 5’ and 3’ RACE products, indicated by black arrows and corresponding to transcripts 2 to 4 or polyadenylation signals A and B, were gel-purified and Sanger sequenced. Other products mentioned in the text are indicated by white arrows.

### MKPV expresses novel “p10” and “p15” accessory proteins

All viable parvoviruses encode NS1 and VP1, and production of these proteins in MKPV-infected tissue was confirmed previously by liquid chromatography-tandem mass spectrometry (LC-MS/MS) (9). However, both MKPV and the extended MuCPV sequence have potential to produce several other polypeptides from ORFs >25 aa in length. We performed a new independent LC-MS/MS analysis of an MKPV-infected kidney and an uninfected kidney, focusing on previously undiscovered MKPV accessory proteins. In addition, we mined our previous LC-MS/MS datasets (PXD010540) (9) for trypsin-derived peptides predicted by these ORFs. These analyses re-identified NS1 and VP1, as expected. They also identified twelve unique peptides (with E-values <0.001) covering 65% of a 14.7 kDa ORF, “p15”, which overlaps the N-terminus of NS1, and two peptides covering the C-terminal 16% of a 9.8 kDa ORF, “p10”, which is situated immediately downstream of the 5’ ITR (see Fig 1B and Table S2 and Table S3). No MKPV-derived peptides were detectable in extracts from uninfected control kidneys. The fact that p15 peptides were more abundant than NS1 peptides in two fully-independent LC-MS/MS analyses suggests that p15 is an abundant protein in MKPV-infected cells, but quantitative assays would be needed to confirm this.

### MKPV mRNA splicing

Previous alignment of Illumina reads with the confirmed MKPV genome indicated the presence of three major introns (9) (see accession MH670587). The two splice donor (SD) and three splice acceptor (SA) sites used by these introns all conform to donor or acceptor consensuses according to a Hidden Markov model for splice site prediction (18). To directly confirm intron splicing, we extracted RNA from kidneys of MKPV^+ve^ *Rag1*^*–/–*^ mice and exposed it to DNase I and Exo I to destroy MKPV DNA. After reverse transcription (RT), we amplified MKPV cDNA using antisense primers 902, 905 or 933 paired with primers 890, 955, 904 or 947, which are mapped in Fig 1A–B. The sense primers “walked” from the hairpin of the 5’ ITR (primer 890) to just upstream of the most 5’ splice donor site (primer 947, Fig 1B). Agarose gel analysis of PCR products (Fig 1C–D) indicated the presence of transcripts 1–4 illustrated in Fig 1B. Splicing of the donor and acceptor sites in transcripts 2–4 was confirmed by Sanger sequencing; similarly, Sanger sequencing confirmed that transcript 1 contained the small 5’ intron intact.

Consistently strong RT-PCR yields using primers 947 and 904 (Fig 1D) suggested that transcription start sites for all four transcripts lie 5’ to MKPV nucleotide 214. In contrast, RT-PCR yields were consistently weak with primer 955, despite this primer yielding strong amplification from MKPV DNA (Fig 1C). Primer 890, located in the hairpin of the 5’ ITR, produced no detectable products in RT-PCR reactions at all, despite being able to produce products from kidney DNA (albeit with relatively low efficiency; Fig 1C). This indicated that MKPV’s major transcription start site(s) lie a short distance downstream of the 5’ ITR.

### Mapping of transcription start and polyadenylation sites by RACE

To precisely map the 5’-ends of MKPV transcripts, we deployed rapid amplification of cDNA ends (RACE) following SMARTer^®^ full-length cDNA synthesis (Fig S1A). Sanger-sequencing of major 5’ RACE products (Fig 1E and Fig S1B) confirmed that they corresponded to transcripts 2– 4 – as labelled in Fig 1E. Two major transcription start sites, P1 and P2, were mapped for transcript 2 (Fig 1A). P1 corresponds to nucleotide 147 with “smearing” to nucleotides 144–146 (transcript 2A), while P2 corresponds to nucleotides 266–267 (transcript 2B). The transcription starts for transcripts 3 to 4 mapped to precisely the same nucleotides as transcript 2A – i.e. P1 (Fig 1B) but not P2. These results were consistent with the RT-PCR reactions in Fig 1D. Since the interior repeat of the MKPV 5’ ITR immediately abuts P1 (Fig 1A), transcription predominantly initiates from very near the 3’-end of the 5’ ITR.

To map polyadenylation sites, we deployed 3’ RACE (Fig 1E, Fig S1A and Fig S1C,). This mapped a major site of polyadenylation (polyA) to nucleotides 3491–2 (“GA”), which lie 13–14 nt 3’ to a polyA signal embedded in the middle of VP1 (signal A in Fig 1B); the “A” lying at nt 3492 prevented single-base precision (see Fig S1C). Polyadenylation sites were also mapped to nucleotides 4292–3 and 4297–8 (both “CA”), which lie 18–24 nt 3’ to the polyA signal at the end of VP1 (signal “B” in Fig 1B). Another polyadenylation site, lying 20–30 nt from the 3’ end of the MKPV genome may also be used (“B3” in Fig 1E), but we did not attempt to map this site precisely because the 3’ ITR is extremely resistant to Sanger sequencing.

We presume that polyadenylation signal A is used by transcripts 1–3 and that signal B is used by transcript 4, but it is possible that transcripts 1–3 use both A and B polyA signals (Fig 1B). Other transcription start sites and polyadenylation sites were indicated by capillary electrophoresis of RACE products (Fig 1E) and by sequencing (Fig S1B–C), but these were very minor. Surprisingly, none of the major 5’ RACE products corresponded to transcript 1, which carries the smallest intron intact (Fig 1D). A very faint 5’ RACE product was detected that might correspond to transcript 1 (“1” in Fig 1E), because it was about 80 bp larger than the 5’ RACE product corresponding to transcript 2A, but it’s yield was too low to be Sanger sequenced.

NS1, which was detected by LC-MS/MS of MKPV-infected kidneys, could theoretically be translated from transcripts 1, 2a or 2b (Fig 1B). The ATG start codon of the p15 ORF, also detected by LC-MS/MS, aligns exactly with the splice acceptor site of transcripts 2a and 2b in Fig 1B (Fig S1). Therefore, p15 could also be produced from transcripts 1, 2a or 2b. In addition, these transcripts potentially encode “NP”, a hypothetical chapparvoviral ORF first noted in a turkey chapparvovirus (13). Transcript 3 encodes a splice variant of NP that we previously dubbed NS2-P (9); however, we have not detected any NS2 or NP peptides by LC-MS/MS so far to confirm production of these polypeptides *in vivo*. Transcript 4 encodes the capsid protein VP1, which was detected by LC-MS/MS (Fig 1B). p10, also detected by LC-MS/MS, is potentially encoded in all transcripts that start from the P1 promoter, but not from transcripts starting at P2.

### MKPV RNA is restricted to the kidney

During the natural course of infection, MKPV DNA was detected in the kidney first, in young adult mice, then appeared in liver, spleen and blood as infection progressed (9). The mapping of MKPV RNA splicing (Fig 1) enabled quantitation of active MKPV infection *via* qPCR for spliced MKPV RNA in different tissue sites. We extracted DNA and RNA from liver, spleen (a proxy for blood) and kidneys of naturally MKPV-infected *Rag1*^*–/–*^ mice, then performed qPCR using DNA or cDNA templates. For DNA, we used NS1 primers 869 and 870, as previously reported (9) (Fig 2A). For cDNA, we used primers 947 and 948 (Fig 2A) and a short extension time, to ensure the product formed from spliced transcripts only. We did not establish a standard curve for cDNA, but instead report the difference between Ct for *Hprt* versus MKPV *Cap* cDNA (Fig 2B). Consistent with our previous study (9), MKPV DNA was much more abundant in kidney than in liver or spleen (Fig 2B). Critically, spliced MKPV RNA was below the detection threshold in liver and spleen, but readily detectable in kidneys (Fig 2B).

**Figure 2.**
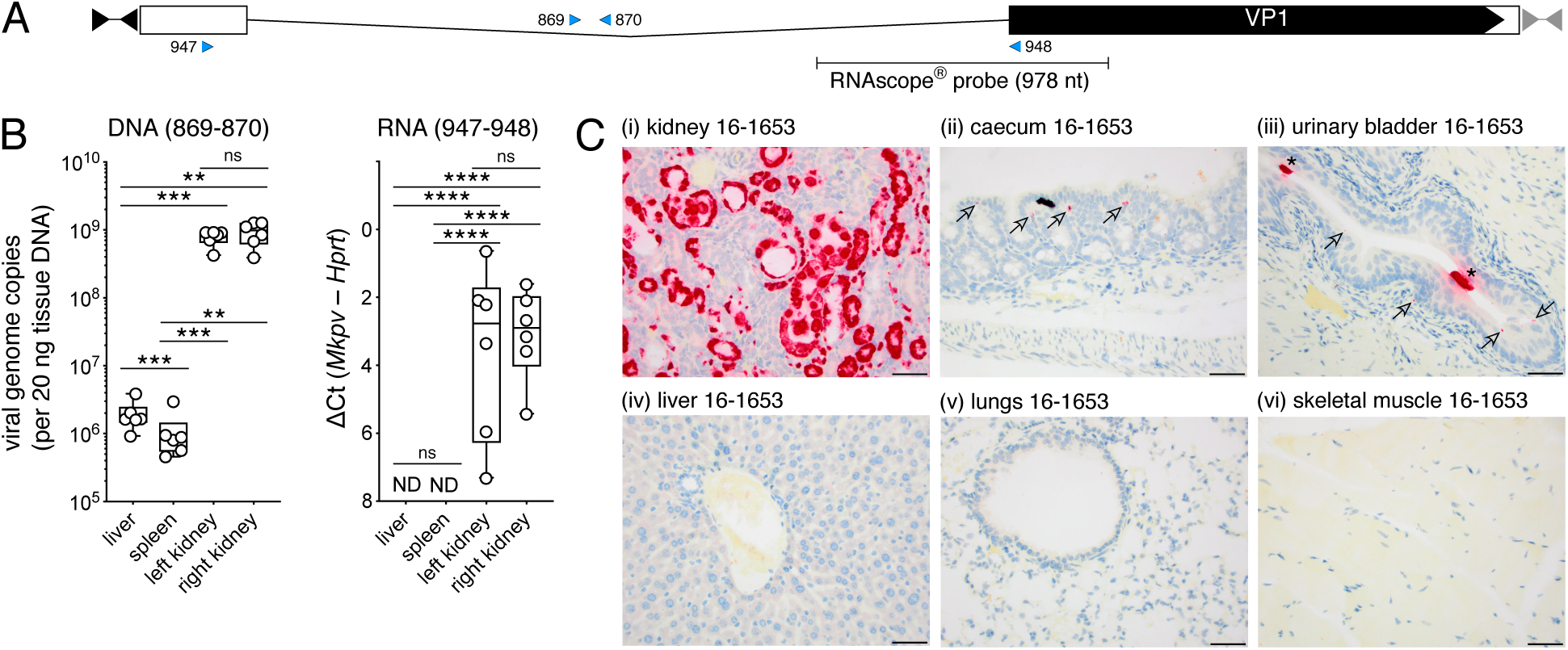
MKPV mRNA is kidney-restricted. **(A)** Placement of PCR primers and the in situ hybridization (ISH) probe relative to the MKPV *cap* transcript (i.e. transcript 4 in Fig 1). **(B)** Quantitation by qPCR of MKPV genomes (left) or MKPV *cap* mRNA (right) in organs of naturally-infected *Rag1*^*– /–*^ mice, using primers 869-870 or 947-948, respectively (Tukey’s box and whisker plots; n = 8). MKPV DNA is presented as viral genome copies. *Cap* mRNA abundance is indicated by Ct relative to RT-qPCR for *Hprt* mRNA. ND = not detected. Significance is indicated by asterisks (*, P<0.05; **, P<0.01, ***, P<0.001, ****, P<0.0001; ns, p>0.05; 1-way paired ANOVA with Tukey’s multiple comparisons test). **(C)** ISH for MKPV nucleic acids in necropsy specimens from NSG mouse 16-1653 housed in MSKCC in 2016. Scale bar = 25 µm. Panel (i) is reproduced from (9). Arrows in panels (ii-iii) indicate mild multi-focal staining in caecum and urinary bladder; asterisks in panel (iii) indicate casts of necrotic tubular cells sloughed from the kidney into the urinary bladder lumen. Full details of ISH outcomes are listed in Table S4.

To examine a greater range of tissues, we deployed an MKPV-specific in situ hybridization (RNAscope®) probe (9) in tissue sections from necropsies of two MKPV^+ve^ NOD-*scid* IL2Rgamma^null^ (NSG) mice. These two mice had histopathologic evidence of chronic inclusion body nephropathy (IBN) and ISH had detected abundant MKPV nucleic acids in tubular epithelial cells (9), as reproduced here (Fig 2Ci). No pathologic change attributed to the virus was observed outside the kidneys on H&E stained sections. Mild multifocal ISH staining was also observed in the caecum mucosal epithelium and lamina propria of one mouse, but not the other, and in the urinary bladder urothelium (mostly umbrella cells) of both mice (Fig 2Cii-iii, arrows). In addition, there were strongly positive cells in the urinary bladder lumen of both mice, which were presumably casts of necrotic tubular cells sloughed from the kidney; an unsurprising finding (Fig 2Ciii, asterisks). No ISH signal for MKPV was detected in any of the 21 other tissues screened (e.g. Fig 2Civ-vi) (Table S4). Another thirteen tissues sampled during necropsy were not probed because the decalcification process used in their preparation for H&E-staining was incompatible with ISH (see Table S4).

Together, these data demonstrate that although MKPV can become widely disseminated during the course of infection, production of MKPV RNA – a definitive marker for active MKPV replication – occurs predominantly, or perhaps exclusively, in the kidneys. We noted previously that MKPV DNA was often more abundant in liver than in other non-kidney sites (9); indeed, MuCPV DNA was originally detected in wild mice in livers and anal swabs (10). Our current analysis revealed significantly higher MKPV DNA levels in liver compared to spleen (Fig 2B; adjusted P = 0.0009), which suggests that the liver may act as an MKPV/MuCPV sink or filter during viremia (19), even though it is not a site of active MKPV replication (see Fig 2Civ). The presence of MKPV DNA in spleen is explained by the onset of viremia, as previously described (9). MuCPV DNA in anal swabs (10) may be related to the faint caecum MKPV ISH signal in Fig 2Cii, or it could reflect contamination of the anal region with urine.

To test whether viruses related to MKPV might also be kidney-restricted, we screened kidneys and livers from seven vampire bats (*Desmodus rotundus*) previously identified as hosts for the chapparvovirus DrPV-1 (14). Kidney DNA from all seven vampire bats was re-confirmed as PCR^+ve^ for DrPV-1 NS1, while liver DNA was uniformly PCR^−ve^ (Fig 3). While this does not prove that DrPV-1 specifically infects kidneys, it demonstrates that DrPV-1 DNA is substantially more abundant in kidneys than livers of host vampire bats.

**Figure 3.**
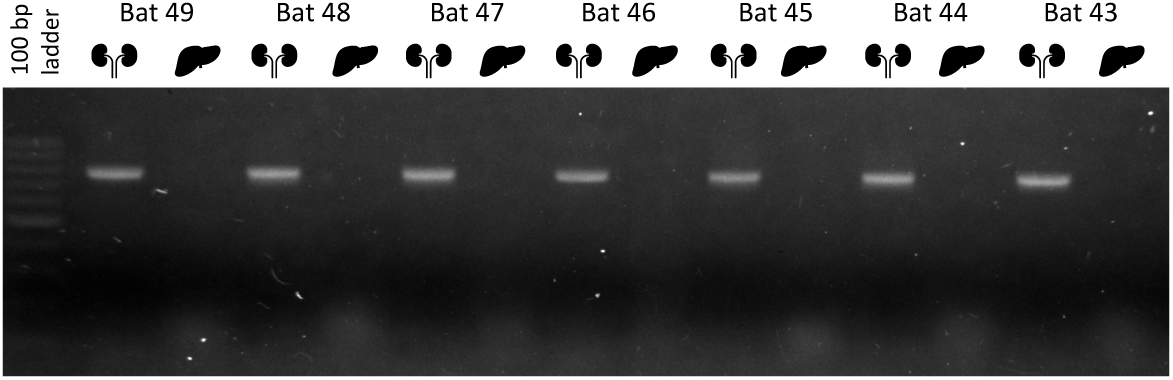
Detection of viral *rep* DNA of DrPV-1 in kidneys and livers of 7 *Desmodus rotundus* vampire bats captured in a rural area of Araçatuba city, São Paulo State, Brazil in 2010 (14). Agarose gel electrophoresis showing 783 bp amplicon specific for the *rep* gene of DrPV-1; M = 100 bp DNA ladder. Sample IDs are shown above and tissues are indicated by silhouettes.

### MKPV geographical distribution and polymorphism

MKPV/MuCPV was reported in five sites previously: in the wild in NYC, USA and in laboratory mice housed in NYC and Baltimore in the USA, and in Sydney plus another Australian city. We screened two additional sets of necropsy specimens from laboratory mice with histologically diagnosed IBN by PCR and detected MKPV DNA in laboratory mice housed at University of North Carolina (Chapel Hill, USA) and in Israel (Fig 4). The specimen from Israel was also probed using ISH and abundant MKPV nucleic acids localised to tubular epithelial cells were detected (Fig 4). This increases the number of sites in which MKPV is associated with mouse kidney disease to six sites in three continents.

**Figure 4.**
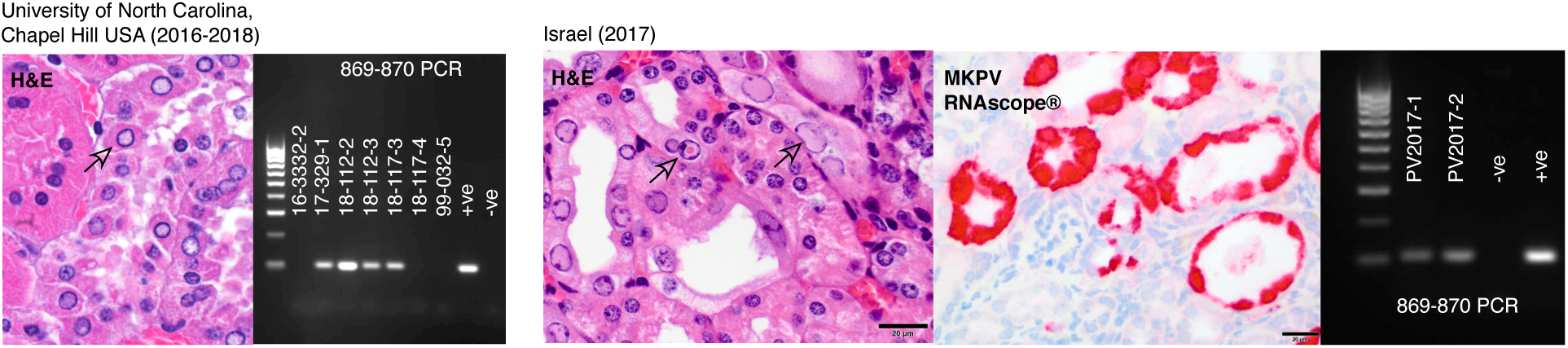
Haematoxylin/eosin (H&E)-staining of historical formalin-fixed paraffin-embedded (FFPE) mouse kidney necropsy samples from the University of North Carolina, Chapel Hill, USA and Israel, paired with agarose gels of 25 cycle 869-870 PCR for MKPV DNA using DNA extracted from FFPE kidney shavings of necropsies from the same sites. For the Israel specimen, RNAscope for MKPV nucleic acids was also performed. Arrows show examples of inclusion bodies in each H&E stain. PCR panels include a 100 bp marker at left and control DNA at right from MKPV-infected Centenary Institute *Rag1*^*–/–*^ mice (+ve) or MKPV-free *Rag1*^*–/–*^ mice (-ve, sourced from Australian BioResources, Mittagong NSW).

In our previously reported investigation of the association of MKPV and IBN in laboratory mice in the facilities of Memorial Sloan Kettering Cancer Center and Weill Cornell Medicine (MSK-WCM) over an 11 year period, the virus was retrospectively detected by 25-cycle conventional PCR in 29 of 34 mice affected by IBN, while it was never found in mice without the disease (9). We attributed the negative results in 5 mice affected by IBN as false negatives due to the limitation of 25-cycle PCR sensitivity on formalin-fixed paraffin-embedded (FFPE) tissues. Since then, we have performed ISH in these 5 cases and detected MKPV nucleic acids in 4 of them (Table S5). The ISH negative mouse was an immunocompetent animal with minimal histological lesions and was co-housed with an ISH positive animal. Therefore, we have demonstrated the presence of MKPV all cases of IBN at MSK-WCM in a period of 11 years.

Alignment of our original CI-MKPV, MSKCC-MKPV and wild NYC MuCPV sequences revealed numerous single nucleotide variations (SNVs) and a two-base insertion in CI-MKPV; Sanger and Illumina sequencing data also revealed a few SNVs within each virus strain (Fig 1A). One notable SNV was the insertion of an extra “C” in the 3’ ITR of a sub-strain present in one CI mouse, converting the sequence C_4_G_4_ to C_5_G_4_ in the interior repeat (“▲” in Fig 1A), without the insertion of a complementary base in the exterior inverted repeat. This SNV creates an extra 1 nt bubble in the structure predicted to be formed by the 3’ ITR (Fig S2), but whether it results in viable virus remains to be determined. A window of concentrated NS1 polymorphisms – boxed in blue in Fig 1A – was selected for sequencing in a larger sample of viruses. This region was amplified (using primers 934 and 935; Fig 1A) from kidney FFPE-specimens or from randomly-selected faecal samples sent to Idexx Laboratories (Columbia, Missouri) from multiple laboratory facilities (in the USA, Canada, Europe and Israel), then Sanger sequenced from both ends. SNVs were collated in a 267 bp window. Including a partial MKPV NS1 sequence identified by whole animal metagenomics of *Mus musculus* living wild in Xinjiang, China (Accession MG679365), this analysis identified 22 MKPV sub-strains, varying by 3–22 SNVs from the consensus sequence (Fig 5). The MSK-WCM colonies provided the largest set of time-shifted samples for the same location, from 2007 to present. There was no clear evidence of one strain replacing another over time in the MSK-WCM colonies. Instead, more than one strain was present in the MSK-WCM colonies at most timepoints, and two sub-strains present in MSK-WCM in 2008–2009 and 2015– 2017 were identical (within our SNV window) to sub-strains from the University of North Carolina (2018) and Johns Hopkins University (2006), respectively. All of the wild NYC samples shared some SNVs with laboratory strains, mostly located in the same continental region. It was notable that the wild NYC sample Q-055 shared four SNVs with Australian laboratory mice (Fig 5), and the wild NYC sample M-118 shared three SNVs with lab specimens from Europe and Israel (Fig 5). This is consistent with MKPV being carried within immune-deficient lab mice when they were live-exported from the USA to labs outside the Americas, but does not prove it.

**Figure 5.**
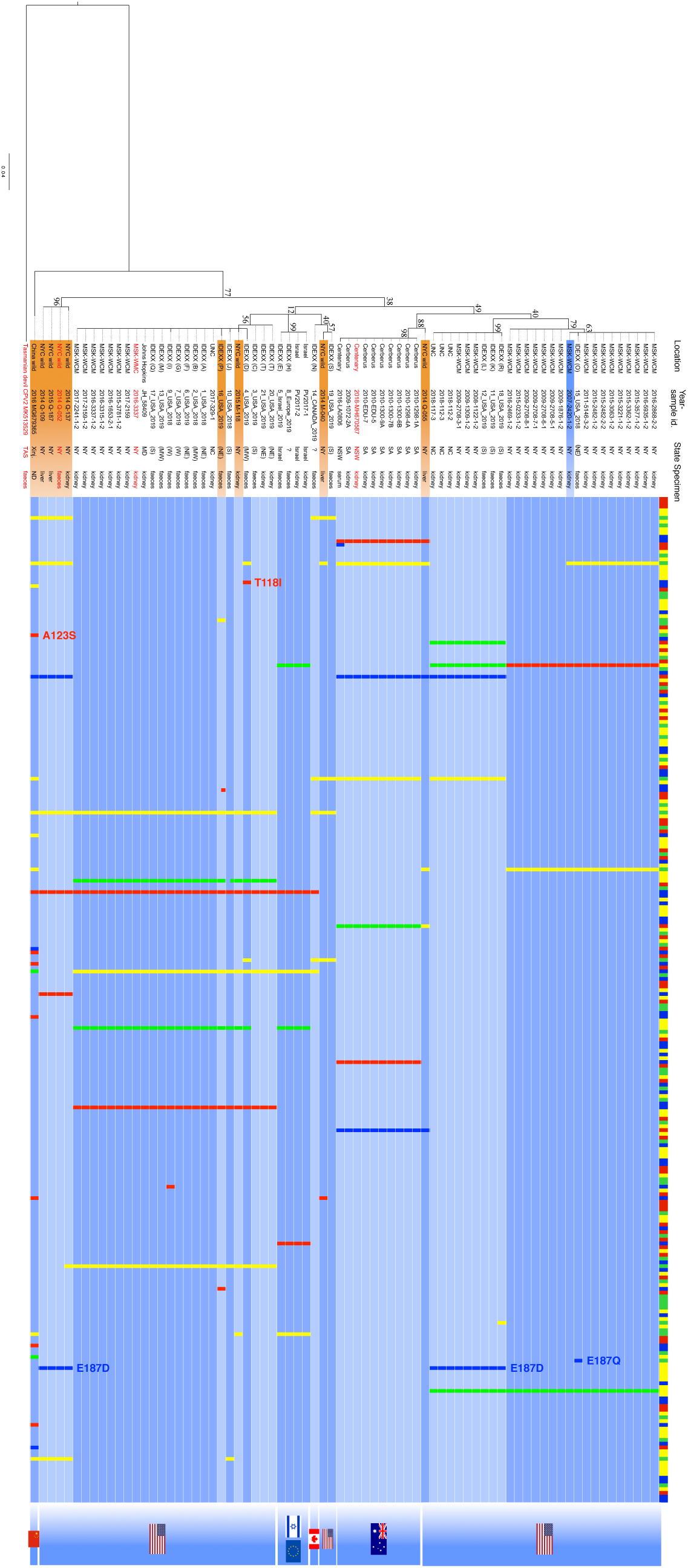
Maximum likelihood phylogenetic tree of MKPV/MuCPV sub-strains based on a 267 bp region in NS1 (see blue box in Fig 1A), using the Tasmanian devil CPV2 sequence (15) as outlier. The scale bar represents units of substitutions per site. Bootstrap nodal support values are indicated. The provenance of each sequence is indicated by text at left and by flags at right, with red text indicating accessions, from top to bottom, MH670587, MH670588, MF175078 and MG679365.1; for IDEXX BioAnalytics pathology samples, donor institutions are identified by an anonymizing code unique to each institution and by a geographical region where known – each in brackets. Shading over the text indicates infection of an immune-competent strain (blue = laboratory mouse, orange = wild-caught mouse). The coloured bar at the top indicates the consensus sequence (yellow = A, green = G, blue = C, red = T). SNVs varying from the consensus are presented as in Fig 1A, with colour-coding to indicate the non-consensus base. Non-synonymous SNVs are indicated by “XnnnX”.

Virtually all SNVs in the 934–935 window were synonymous, with just four exceptions (Fig 5): Glu187Asp present in 2008-9 MSKCC, 2014-2015 wild NYC and 2018-2019 samples from UNC and another nearby location; Glu187Gln in a 2018 sample from the USA Northeast, Thr118Ile in a 2019 sample from the USA Midwest; and Ala123Ser in wild mice from Xinjiang province in China. None of these non-synonymous mutations are likely to affect NS1’s tripartite helicase domain.

### MKPV prevalence

To estimate the prevalence of MKPV in research mice, mouse faecal samples that were submitted to IDEXX BioAnalytics over a seven-month period and representing 78 biomedical research institutions were tested for MKPV by qPCR. Overall prevalence was 5.1%, with 178 positive samples out of 3,517 samples tested. Immune status is unknown for most of the samples. Of those samples designated as representing immunodeficient mice, 16 were positive out of 171 tested (9.4%), and for samples designated as representing immunocompetent mice, 56 out of 513 (10.9%) tested positive for MKPV. It should be noted that many of the faecal samples are likely to represent soiled-bedding sentinel mice, and the prevalence among sentinel mice may differ from colony animals, which may be immunocompetent or immunodeficient, based on a variety of factors including shedding dynamics and efficiency of transmission by soiled bedding.

Faecal samples from multiple time-points from a single pet shop in Columbia, MO were also submitted to IDEXX BioAnalytics. Seventeen samples were qPCR-positive out of 73 tested (23.3%). In toto, the data establish that MKPV-infection is common in wild, laboratory and pet mice globally.

### The assembly of a closely-related chapparvovirus sequence from primate kidney DNA

In addition to MKPV/MuCPV, metagenomics or mining of draft genome assemblies has assembled several near-complete chapparvoviral genomes from five mammals (vampire bat (14), fruit bat (20), rat (21), pig (11) and the marsupial *Sarcophilus harrisii* (Tasmanian devil) (15), three birds (chicken (22), brown mesite (14) and red-crowned crane (23)), and a reptile (the pit viper (14)). Partial genomes lacking substantial 5’- or 3’-coding sequences have also been assembled from numerous species, including from the draft genome of the capuchin monkey (*Cebus capucinus imitator*) (14). The draft capuchin genome was assembled using DNA extracted from the kidney (24). We used the genome of MKPV as a scaffold to re-arrange two sequence fragments present in *Cebus imitator* scaffold NW_016109986 into a near-complete parvoviral genome, but lacking ITRs and with a probable gap in NS1 (Fig S3). Using SAMtools (25), we mapped high quality reads in the complete capuchin kidney NGS dataset (24) to this draft viral genome, which in-filled a 5 nt gap in the NS1 sequence compared to scaffold NW_016109986.1. By recovering sequences from “soft”-clipped reads at the 5’- and 3’-ends of this new alignment (Fig S3), we produced an extended but still incomplete draft viral genome. A final round of read assembly and recovery of clipped ITR sequences produced a complete parvovirus genome with apparently full-length ITRs. We have dubbed this complete viral genome “capuchin kidney parvovirus” (CKPV, Accession MN265364). The kidney DNA sample used to produce CKPV (and the draft *Cebus imitator* genome) was extracted in a biological safety cabinet inside a BSL-2 laboratory facility with strict CL3 protocols to minimise contamination with foreign DNA, but it is no longer available, so we cannot strictly rule out the possibility that CKPV is an extraneous contaminant.

The CKPV genome is strikingly similar to MKPV (Fig 6A) and encodes proteins homologous to MKPV p15 and p10; the VP1, NS1, p15 and p10 proteins of CKPV are 77%, 71%, 76% and 55% identical to their MKPV counterparts, respectively (Fig 6B). Furthermore, the U2-dependent splice donor and acceptor sites used in MKPV for expression of VP1 and for the hypothetical accessory protein NS2 (9) are conserved in CKPV (Fig 6A and Fig S4). We predict a spliced NS2 protein in CKPV that is a remarkable 84% identical to MKPV’s spliced NS2 protein (Fig 6B). In contrast, non-spliced variants of NS2 (“NP-L, −I and −F”) are less conserved because CKPV lacks the in-frame ATG codons upstream of NS2 exon 2 that are present in MKPV (Fig 6A). Finally, the hairpin structures with the lowest Gibbs-free energy predicted for minus strand CKPV ITRs are strikingly similar to the structures predicted for minus strand MKPV ITRs (Fig 6A).

**Figure 6.**
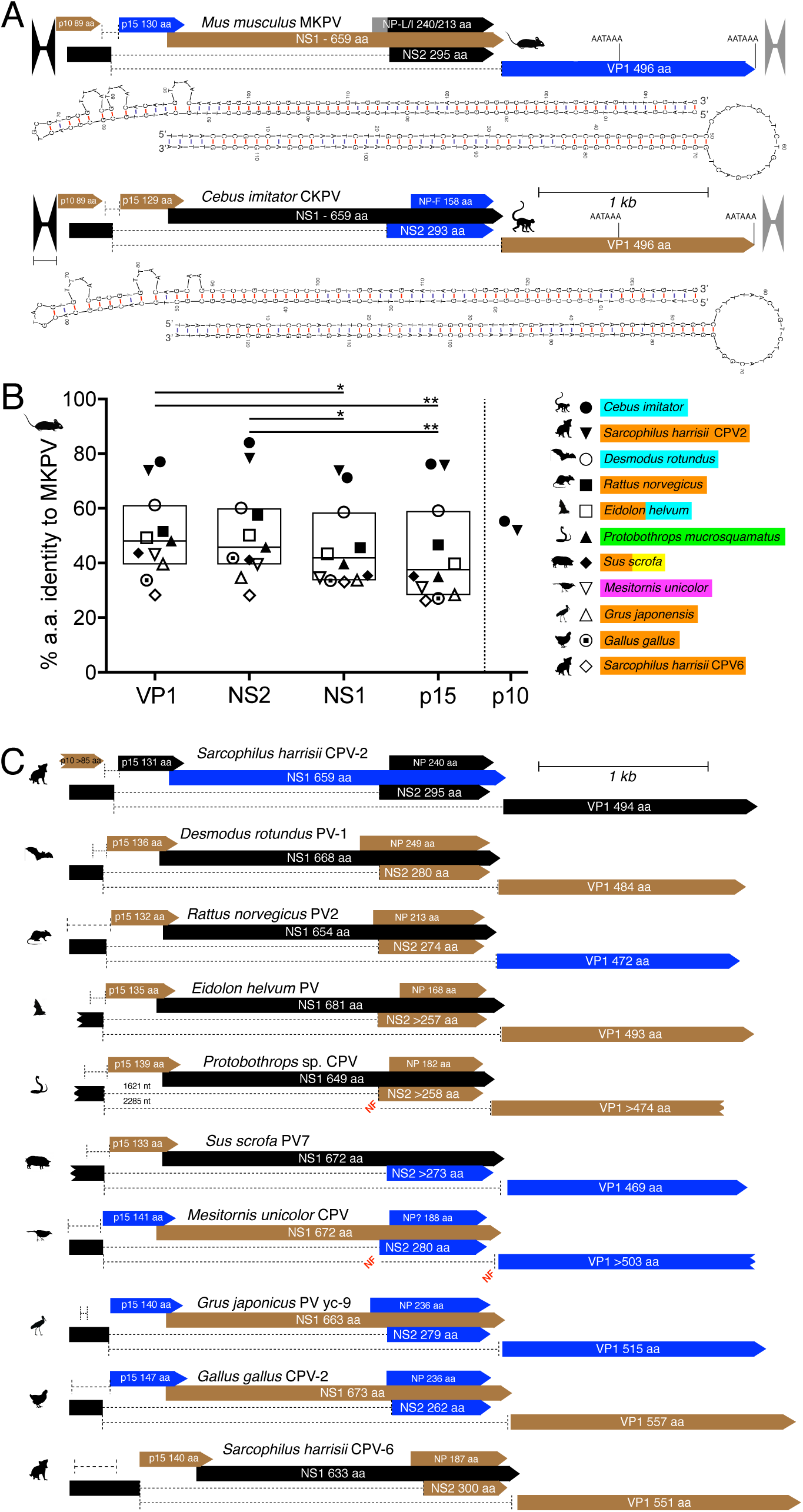
VP1, NS1, NS2 and p15 ORFs, introns and ITR structures are conserved in tetrapod chapparvoviruses. **(A)** Maps comparing the complete genomes of MKPV (top) and CKPV (bottom), to scale. “Bowties” indicate ITRs. Three colours are used to indicate the three plus-strand reading frames. Dashed lines indicate introns – actual (MKPV) or hypothetical (CKPV). The lowest Gibb’s energy structures predicted for the minus-strand ITR regions (36) are shown below each map, with the 5’ ITR above the 3’ ITR. **(B)** The percentage identity at the amino acid level between MKPV ORFs and the corresponding ORFs from the nine other near-complete tetrapod chapparvoviral genomes currently available (MUSCLE alignment (37)). Tukey’s box and whiskers are used. Significant differences (non-parametric Friedman test) are indicated as in the Fig 2 legend. Colours indicate source(s) of the virus sequences: blue – kidney or urine, orange – faeces, yellow – lungs or respiratory tract, green – liver, pink – muscle. **(C)** Maps (same scale, colours and symbols as panel A) indicating conserved p10, p15, NS2, NS1 and VP1 ORFs and putative introns in ten other tetrapod chapparvoviral genomes. The sources for the genomes are: (11, 14, 15, 20-23). Ragged ends indicate incomplete ORFs that continue beyond the currently available sequence. Slice donor and acceptor sites for all introns shown in A and C are listed in Fig S4.

### ORFs encoding p15 and spliced NS2 are conserved in all tetrapod chapparvoviruses

Encouraged by ORF conservation between MKPV and CKPV, we searched all near-complete tetrapod-hosted chapparvovirus genomes for conserved ORFs using “Genie” software (18) to identify likely splice donor and acceptor sites for these ORFs. In all cases, we found a p15-like ORF and a 2-exon NS2-like ORF (Fig 6C and Fig S4). Furthermore, consensus U2-dependent splice donor and acceptor sites were predicted to produce transcripts equivalent to MKPV transcripts 1–4 in nearly all cases (Fig 6C and Fig S4). The exceptions (annotated as “NF” for “not found” in Fig 6C) were that Genie software did not find splice acceptor sites upstream of the *Protobothrops* sp. or *M. unicolor* virus VP1 regions, nor for the *M. unicolor* virus NS2 exon 2. Nonetheless, manual alignment of reading frames implies a functional acceptor site for the *M. unicolor* virus NS2 exon 2 and “acceptor-like” sites a short distance upstream of VP1 in the *Protobothrops* sp. and *M. unicolor* viruses (see Fig S4).

Based on our parsing of chapparvovirus genomes, we produced consensus protein sequences for p15 and NS2 using T-Coffee (26) and searched Pfam and SwissProt for proteins carrying similar motifs on the HMMER server (see Methods), but did not find any significant matches. Nonetheless, p15 universally carries clusters of basic amino acids in the C-terminal region that are suggestive of nuclear localisation signals (NLS).

While all tetrapod chapparvoviruses we have examined potentially express a two-exon NS2 protein, not all of them carry ATG codons upstream of, or within, NS2 exon 2 necessary for expression of the shorter single exon variant of NS2 originally annotated as the hypothetical NP polypeptide in turkey chapparvovirus (13) – see *Sus scrofa* PV7 (Fig 6C). This suggests that expression of two-exon NS2 is preferred over single-exon NP. However, some parvoviruses are known to use cryptic translation start codons (27), so all tetrapod chapparvoviruses might nonetheless express NP in addition to, or instead of, spliced NS2.

p15 was significantly less conserved across tetrapod chapparvoviruses compared to VP1 and spliced NS2 polypeptides (Fig 6B). On the other hand, VP1 and NS2 were significantly more conserved than NS1 (Fig 6B). To the limited extent that parvovirus tropism may be dictated by the capsid (8), our identification of VP1 as the most conserved ORF suggests that chapparvoviruses might target a similar cell type in all tetrapod hosts. Similarly, spliced ORF conservation suggests that NS2 plays an important role in the virus life cycle even though we have not been able to directly detect NS2 protein in MKPV-infected tissue to date.

### The p10 ORF is not present in most chapparvoviruses

Apart from the two closest MKPV relatives, which were found in a capuchin monkey and in faeces of two Tasmanian devils (Fig 6B), an ORF corresponding to p10 was not identifiable in any other chapparvoviruses we examined (Fig 6C), even when complete NS2 exon 1 sequence was available (e.g. in DrPV-1). The capuchin CKPV and Tasmanian devil CPV-2 viruses encode VP1, NS2, NS1 and p15 proteins with >71% amino acid identity to the MKPV proteins, but their p10 proteins are <60% identical to MKPV p10 (Fig 6B). Thus, p10 is poorly conserved, even absent in many chapparvovirus species, unlike the accessory proteins p15 and NS2. Therefore, p10 would appear to be less important than p15 and NS2 in chapparvoviral life cycles.

## Discussion

Our data add to the association between MKPV and chronic IBN in immune-deficient laboratory mice and demonstrate that MKPV is distributed worldwide. MKPV can be detected in immune-sufficient laboratory mice (9) as well as in wild-living mice in the USA (10) and China, which indicates that MKPV does not require immune-deficiency to propagate and that mice are a natural MKPV host. The virtual absence of SNV diversity in MKPV samples from Australian laboratory mice spanning a decade (Fig 5) suggests that a single MKPV strain was imported into Australia and has been transmitted horizontally in laboratory mice since, with little or no re-infection from wild mouse sources. In contrast, infection of mouse colonies in the USA by virus from wild mouse pools appears to have occurred repeatedly, but these infections may have occurred prior to the establishment of modern barrier facilities. The MKPV-infected Australian colonies we first reported (9) were all descended from the *Rag1*^tm1Bal^ strain (28) – imported into Australia from northeast USA in the mid-1990s. Fig 5 indicates that this strain is the original source of MKPV in Australian laboratory mice, because all MKPV^+ve^ Australian lab mice carry the same MKPV strain. Shared SNVs between the Australian MKPV strain and wild mouse sample Q-055 (Fig 5) suggest, but do not prove, that infection originally occurred in the USA. Our MKPV-free *Rag1*^*–/–*^ colony descends from the *Rag1*^tm1Mom^ strain (29) supplied by The Jackson Laboratory (Bar Harbor, Maine, USA). *Mus musculus* is not native to Australia and is thought to have arrived onboard European ships 230 to 420 years ago (30). Given that that other parvoviruses are prevalent in feral house mice in Australia (31) and a virus closely related to MKPV was found in faeces from Tasmania (15), it appears likely that MKPV will be found in *Mus musculus* living wild in Australia. In contrast to *Mus musculus, Pseudomys* and other native mice have lived in Australia for four million years or more (32). Since *Sarcophilus harrisii* CPV2 is only distantly related to the other CPVs found in Tasmanian devil faeces ((15) e.g. *Sarcophilus harrisii* CPV6, see Fig 6), we speculate that *Sarcophilus harrisii* CPV2 infects native rodents that form part of the diet of Tasmanian devils. Our original proteomic analysis of proteins produced by MKPV (9) was limited to NS1, NS2/NP and VP1. Inclusion of all MKPV ORFs ≥25 amino acids in the current analysis identified MKPV p15 and p10 in two independent MKPV-infected kidneys. The so-far universal presence of ORFs encoding p15 and NS2 in tetrapod-hosted chapparvoviruses demonstrates that these are accessory proteins likely to be important in the life cycle of all tetrapod-adapted *Chapparvovirinae*. In contrast, the hypothetical NP protein and the p10 accessory protein seem to be much less conserved or absent in many chapparvoviruses; this is markedly the case for p10. Neither NS2 nor p15 bear any significant structural homologies to other proteins, so we have no clues to their functions at present.

We have now demonstrated conclusively that MKPV replicates preferentially in the mouse kidney (Fig 2). Furthermore, the absence of any obvious co-replicating virus in MKPV-infected kidneys (9) strongly implies, but does not prove, that MKPV replicates autonomously. Strong conservation of VP1 sequence, accessory proteins (excepting p10) and splicing, plus near-complete kidney-tropism in MKPV combined with at least some degree of kidney-tropism in CKPV and DrPV-1, suggests that viruses closely-related to MKPV are also adapted to kidney niches in distantly-related mammalian hosts, including non-human primates. It is tempting therefore to speculate that many tetrapod-adapted *Chapparvovirinae* preferentially infect the kidneys. The structural conservation between MKPV and CKPV of heterotelomeric ITRs furthermore provides a prototypical genome, illustrated in Fig 6A, for a genus we propose be dubbed *Nephroparvovirus*. The “*Nephroparvovirus”* genome is clearly distinct from the prototypical genomes of *Protoparvovirus, Ambidensovirus, Erythroparvovirus, Bocaparvovirus* and *Dependoparvovirus* (reviewed in (1)).

MKPV infection in *Rag1*^*–/–*^ mice shares clinico-pathological features with polyomavirus-associated nephropathy (PVAN), which is a significant complication in immune-suppressed kidney transplant recipients (9, 33). Our assembly of the complete CKPV genome from the kidney DNA of a capuchin monkey increases the possibility that a *Chapparvovirinae* species might infect human kidneys. Reasoning that urine from immune-suppressed kidney transplant patients is the most likely material in which human chapparvoviral infection might be detected, we mined the published fastq files produced by deep-sequencing the urinary DNA of 27 kidney transplant patients ((34), NCBI accession PRJEB28510), searching for chapparvoviral sequences. However, we found none within the datasets. Indeed, we found no parvoviral sequences of any sort within the datasets, as originally reported (34). This limited sample suggests that human kidney chapparvoviral infection is not widespread – at least not in the USA, but it will nonetheless be worthwhile to determine if antibodies against chapparvoviral VP1 antigens are present in human populations. If antibodies are absent, then recombinant AAV vectors pseudo-typed with chapparvoviral VP1 capsid may be better able to evade antibody-mediated immunity than AAV vectors presently used in the clinic, which are unsuitable for use in the >70% of the human population with pre-existing anti-AAV immunity (35).

## Acknowledgments

We thank G. Duytschaever (University of Calgary), Erica K. Creighton and David C. Eckhoff (IDEXX BioAnalytics), and the staff at the Centenary Institute animal facility for their assistance. Supported by the Australian National Health and Medical Research Council (W.W., B.R., P.B. & J.J.-L.W), the Cancer Institute NSW (B.R. & J.J.-L.W), the Hillcrest Foundation (C.J.J.), the Alfred P. Sloan Foundation (S.H.W.), the National Institutes of Health (U19AI109761 Center for Research in Diagnostics and Discovery, S.H.W.), the National Cancer Institute Cancer Center Support Grant P30 CA008748 (S.M.), the Fundação de Amparo à Pesquisa do Estado de São Paulo, Brazil (No. 17/13981-0 and 18/09383-3, W.M.S, & M.J.F.), the National Sciences and Engineering Research Council of Canada (A.D.M), the Canada Research Chairs program (A.D.M.), the Alberta Children’s Hospital Research Institute (A.D.M. & J.D.O.) and the Beatriu de Pinós postdoctoral programme of the Government of Catalonia’s Secretariat for Universities and Research of the Ministry of Economy and Knowledge (J.D.O.).

## Materials and Methods

### Mice and mouse specimens

A colony of naturally MKPV-infected *Cxcr6*^gfp/gfp^ *Rag1*^*–/–*^ mice (C57BL/6.Cg) was maintained in the Centenary Institute mouse facility (Sydney, NSW, Australia), as previously described (9); MKPV-free *Rag1*^*–/–*^ mice (C57BL/6.Cg), originally sourced from The Jackson Laboratory, were purchased from Australian BioResources (Moss Vale, NSW, Australia). Fresh tissue specimens were harvested from mice immediately after humane euthanasia with approval for mouse care and experimental procedures by the Animal Welfare Committee, Royal Prince Alfred Hospital (Sydney, NSW, Australia; approval number 2017-043) and in accordance with NSW and Australian Federal legislation and the *Australian code for the care and use of animals for scientific purposes* (38). Multiple colonies representing various strains of mice, including NOD.Cg-*Prkdc*^*scid*^ *Il2rg*^*tm1Wjl*^/SzJ (NSG) and Tac:SW (Swiss Webster) were maintained at Memorial Sloan Kettering Cancer Center (MSK) and Weill Cornell Medicine (WCM), as described previously (9). Mouse care and experimental procedures were approved by the MSK and WCM Institutional Animal Care and Use Committee and maintained in accordance with the National Academy of Sciences’ Guide for the Care and Use of Laboratory Animals in AAALAC International-accredited facilities.

Samples analysed for MKPV nucleic acids by PCR or ISH (RNAscope) were formalin-fixed paraffin-embedded (FFPE) specimens archived from historical necropsies of diseased mice from these or other colonies, as described previously (9). DNA or total RNA was extracted from fresh tissue or FFPE samples using QIAamp^®^ DNA Mini Kits or RNeasy® Mini Kits from Qiagen (Hilden, Germany), according to the manufacturer’s instructions – with modifications as described previously (9). Total nucleic acids were extracted from mouse samples submitted to IDEXX BioAnalytics using a commercially available platform (NucleoMag® VET Kit; Macherey-Nagel GmbH & Co. KG, Düren, Germany).

### PCR from MKPV DNA and phylogenetic analysis

All primers mentioned in this report are listed in Table S1. Most PCRs used 100 ng input DNA, Phire II hot start Mastermix (Thermo Fisher, Vilnius, Lithuania) and 0.5 µM primer pairs. Cycling conditions were as follows: initial denaturation at 98°C for 30 s, followed by 30 cycles (Fig 1C and Fig 5) or 25 cycles (Fig 4) of denaturation at 98°C for 5 s, annealing at 58°C (Fig 1C and Fig 5) or 48°C (Fig 4) for 5 s, and extension at 72°C for 15 s, and concluded with a final extension at 72°C for 5 min. 5 or 7 μL of the completed PCR product was then loaded onto 1.5% agarose (Vivantis, Selangor Darul Ehsan, Malaysia) gels prepared in 1X TAE buffer (Invitrogen, Grand Island, NY, USA) and 1:10,000 dilution of GelRed (Biotium, Fremont, CA, USA). Electrophoresis was conducted at 110 A for 60 min before imaging with G:BOX (SynGene, Cambridge, UK) using Syngene’s GeneSnap v7.05 software. PCR amplifications of sequences missing from the 5’- and 3’ends of the previously-described NYC and MSKCC strains of MuCPV/MKPV used primers 890 and 889 for the 5’end or 891 and 893 for the 3’-end, using 30 to 35 cycles of PCR as above. The products were then Sanger-sequenced (Macrogen, Seoul, South Korea).

PCR amplifications for MKPV SNVs used primers 934 and 935 (Table S1) and were mostly perfomed using Phire II hotstart Mastermix as described above, with the following exceptions. PCR for SNVs at IDEXX BioAnalytics used LA Taq™ (TaKaRa Bio, Ōtsu, Japan) and 20 µM primer pairs. Cycling conditions were as follows: initial denaturation at 94°C for 1 min, followed by 40 cycles of denaturation at 94°C for 30 s, annealing at 58°C for 30 s, and extension at 72°C for 30 s, and concluded with a final extension at 72°C for 5 min. PCR for SNVs from wild NYC mice used AmpliTaq Gold 360 Master Mix (Applied Biosystems, Foster City, CA), 50 µM primers and cycling conditions as follows: initial denaturation for 95°C for 8 min, followed by 10 cycles of denaturation at 95°C for 30 s, annealing at 60°C (decreasing by 0.5°C per cycle) for 30 s, and extension at 72°C for 30 s, then a further 35 cycles with similar conditions aside from an annealing temperature of 55°, and concluded with a final extension at 72°C for 7 min. Both strands of PCR products were Sanger sequenced at Macrogen (S Korea; Australian specimens), or at Genewiz Inc. (South Plainfield, NJ; IDEXX and wild NYC specimens). For phylogenetic analysis, nucleotide sequences were aligned in Geneious 10.2.3 (39), and exported to MEGA6 (40) where model selection was performed. A maximum likelihood tree was constructed using the Tamura 3-parameter model (41) with 1000 bootstrap repetitions. The newick tree was exported to FigTree (v1.2.2, http://tree.bio.ed.ac.uk/software/figtree/) for annotation.

### Testing for prevalence of MKPV DNA by IDEXX BioAnalytics

MKPV detection by IDEXX BioAnalytics used a real-time PCR assay based on the IDEXX Laboratories, Inc. proprietary service platform. The MKPV real-time PCR primers and hydrolysis probe were designed with PrimerExpress® version 3.0 (Applied BioAnalytics™; Waltham, MA, USA) using the genome sequence available in GenBank. The assay was designed and validated to detect 1-10 template copies. Analysis was performed at IDEXX BioAnalytics (Columbia, MO, USA) with standard primer and probe concentrations using the master mix LightCycler® 480 Probes Master (Roche Applied Science, Indianapolis, IN, USA) in a commercially available instrument (LightCycler® 480; Roche Applied Science). In addition to positive and negative assay controls, a hydrolysis probe-based real-time PCR assay targeting universal prokaryotic (16s rRNA) and eukaryotic reference genes (18s rRNA) was amplified for all samples to confirm the presence of amplifiable DNA and absence of PCR inhibition.

### PCR, RACE and qPCR from MKPV RNA

For splice site confirmations, MKPV-infected kidney RNA was treated or mock-treated with Turbo™DNase (Thermo Fisher, 0.4 U/µg RNA) and ExoI (New England Biolabs, 2 U/µg RNA) for 30 min at 37°C, followed by incubation with DNase Inactivation Reagent (Thermo Fisher, 0.2ul/ µg RNA). First-strand cDNA was produced using random hexamer (Bioline, London, UK) priming and SuperScript III reverse transcriptase (Thermo Fisher, Carlsbad, CA, USA) according to the manufacturer’s protocol. PCR was then performed using input cDNA equivalent to 4 ng RNA and Phire II hot start Mastermix (Thermo Fisher, Vilnius, Lithuania), exactly as described for Fig 1C in the preceding section.

Rapid amplification of cDNA ends (RACE) was performed using reagents from an In-Fusion® SMARTer® Directional cDNA Library Construction Kit (TaKaRa) and custom primers. First strand cDNA synthesis was performed using the SMARTer CDS primer (Table S1), SMARTScribe reverse transcriptase (RT) and 500 ng of MKPV-infected kidney RNA, according to the kit instructions. A switch to using a “Template Switch” oligonucleotide (Table S1) as template occurred during first strand synthesis when the RT incorporated untemplated dCMP nucleotides after encountering the 5’ end of an mRNA template (see Fig S1). For 5’-RACE, a 2 µL aliquot of the first strand cDNA diluted 1:50 was amplified using Phire II hot start Mastermix (ThermoFisher, Vilnius, Lithuania) and 0.12 µM of 5’-RACE primer (Table S1) paired with an MKPV-specific antisense oligodeoxynucleotide as return primer. For MKPV transcripts 2a and 2b, primary 5’-RACE with MKPV primer 902 produced two dominant products after 34 PCR cycles (see Fig 1E). For transcripts 3 or 4, primary 5’-RACE used MKPV-specific return primers 905 or 933, respectively, for 25 PCR cycles; a 2 µL aliquot of primary PCR reaction diluted 1:100 was then used as template for semi-nested secondary 5’-RACE using MKPV-specific return primers 900 or 948, respectively, for 34 PCR cycles to produce a single dominant product in each case (see Fig 1E). For 3’-RACE, a 2 µL aliquot of the first strand cDNA diluted 1:50 was amplified in a similar way, using an MKPV-specific sense oligodeoxynucleotide paired with the 3’ RACE primer (Table S1) as return primer. For polyadenylation site A, primary 3’-RACE used MKPV-specific primer 932 and produced a dominant product after 34 PCR cycles (see Fig 1E). For polyadenylation site B, primary 3’-RACE was performed using MKPV-specific primer 940 for 25 cycles; a 2 µL aliquot of the primary 3’-RACE reaction diluted 1:100 was then used as template for semi-nested 3’RACE using MKPV-specific primer 891 for 34 PCR cycles to produce two or three dominant products (see Fig 1E). Sizes and yields of RACE products (2 µL aliquots) were determined using a Fragment Analyzer (Advanced Analytics; now Agilent, Santa Clara USA) equipped with 55 cm electrophoresis capillaries and reagents capable of resolving dsDNA fragments between 35 and 1500 bp (see Fig 1E), according to the manufacturer’s procedures. Signal traces from each capillary were converted into pseudo-gel images using PROSize 2.0 software (Agilent Technologies, Santa Clara, CA, USA – see Fig 1E). The remaining bulk of the RACE reactions were then resolved by conventional 1.5% agarose gel electrophoresis, as described (9). DNA bands stained with GelRed (Biotium, Freemont CA, USA) were excised under blue light. The DNA was extracted from the excised agarose using a PCR product purification kit (Promega, Madison, Wisconsin, USA), following the manufacturer’s instructions for agarose-embedded DNA, then Sanger-sequenced (Macrogen, Seoul, S Korea) using the appropriate MKPV-specific primer.

qPCR for MKPV DNA was performed exactly as described before (9), using primers 869 and 870, an annealing temperature of 48°C and an extension time of 0.5 min. For RNA qPCR, RNA equivalent to 1 ug tissue was pre-treated with DNase, as described above, then reverse-transcribed using oligo-dT primer and SuperScript III reverse transcriptase (ThermoFisher, Carlsbad, CA, USA) at 50 °C for 60 min. First-strand cDNA equivalent to 40 ng tissue was then used as template for qPCR reactions using primers 947 and 948, iQTM SYBR® Green Supermix (Bio-Rad, Hercules, CA, USA), an annealing temperature of 68.5°C and an extension time of 0.5 min.

### *Cebus imitator* genome assembly

The *Cebus capucinus imitator* genome was assembled as described (24). In brief, DNA for shotgun sequencing was derived from the kidney of an adult male (id no. Cc_AM_T3) that was killed by a vehicle in Costa Rica. Samples were transported to the laboratory of A.D.M. at Washington University in St Louis, MO under CITES export permit 2015-CR1258/SJ (no. S 1320). Total sequence genome input coverage on the Illumina HiSeq 2500 instrument was approximately 81x (50x fragments, 26x 3kbs, and 5x 8kbs) using a genome size estimate of 3.0Gb. The combined sequence reads were assembled using ALLPATHS-LG software (42) to produce assembly GCA_001604975.1.

### *Desmodus rotundus* specimens and DrPV-1 PCR

DNA was extracted from kidney and liver samples previously obtained from seven DrPV-1^+ve^ *Desmodus rotundus* individuals (vampire bats) captured in a rural area of Araçatuba city, São Paulo State, Brazil, in June 2010 (14) using a QIAamp DNA Mini Kit (Qiagen, USA). Sample collection and handling procedures were approved by the Brazilian Committee on Animal Experimentation (protocol number 00858-2012) and Chico Mendes Institute for the Conservation of Biodiversity; protocol numbers 12.751-3/2009 and 27.346-1/2011. DNA was screened for DrPV-1 sequences using Platinum® Taq DNA polymerase and high fidelity PCR buffer (Invitrogen), primers Chap-DRPv-fwd and Chap-DRPv-rev, an initial temperature of 94°C for 0.5 min, and 35 cycles of 94°C for 0.25 min, 57°C for 0.5 min and 68°C for 1 min. Products were resolved by agarose gel electrophoresis.

### Tissue staining

Tissue sections were prepared and stained with haematoxylin and eosin (H&E) or with an ISH (RNAscope) probe specific for MKPV nucleic acids and positive and negative control probes as previously described (9). Positive results on caecum and urinary bladder were confirmed by duplicate staining on two serial sections performed in two independent staining runs.

### LC-MS/MS

Kidney protein extracts from an age- and gender-matched pair of MKPV-infected and uninfected *Rag1*^*–/–*^ mice were prepared and digested with trypsin as described (9). Using an Acquity M-class nanoLC system (Waters, USA), 5 µL of each sample was loaded at 15µL/min for 3 minutes onto a nanoEase Symmetry C18 trapping column (180 µm × 20 mm) before being washed onto a PicoFrit column (75 µmID × 300 mm; New Objective, Woburn, MA) packed with Magic C18AQ resin (3 µm, Michrom Bioresources, Auburn, CA). Peptides were eluted from the column and into the source of a Q Exactive Plus mass spectrometer (Thermo Scientific) using the following program: 5-30% MS buffer B (98% Acetonitrile + 0.2% Formic Acid) over 90 minutes, 30–80% MS buffer B over 3 minutes, 80% MS buffer B for 2 minutes, 80–5% for 3 min. The eluting peptides were ionised at 2400V. A Data Dependant MS/MS (dd-MS^2^) experiment was performed, with a survey scan of 350–1500 Da performed at 70,000 resolution for peptides of charge state 2+ or higher with an AGC target of 3e6 and maximum Injection Time of 50ms. The Top 12 peptides were selected fragmented in the HCD cell using an isolation window of 1.4 m/z, an AGC target of 1e5 and maximum injection time of 100ms. Fragments were scanned in the Orbitrap analyser at 17,500 resolution and the product ion fragment masses measured over a mass range of 120-2000 Da. The mass of the precursor peptide was then excluded for 30 seconds.

The MS/MS data files were searched using Peaks Studio X against a database comprised of the *Mus musculus* proteome (UniProt UP000000589) plus all MKPV ORFs >25 amino acids plus a database of common contaminants; with the following parameter settings. Fixed modifications: none. Variable modifications: propionamide, oxidised methionine, deamidated asparagine. Enzyme: semi-trypsin. Number of allowed missed cleavages: 3. Peptide mass tolerance: 10 ppm. MS/MS mass tolerance: 0.05 Da. The results of the search were then filtered to include peptides with a –log_10_P score that was determined by the False Discovery Rate (FDR) of <1%, the score being that where decoy database search matches were <1% of the total matches.

### Bioinformatics

Potential splice donor and acceptor sites in chapparvoviral genomes were sought using “Genie” software (18) online (http://www.fruitfly.org/seq_tools/splice.html). Alignments of chapparvoviral proteins p15, p10, NS1, NS2, NP and VP1 were performed by MUSCLE (37) or T-Coffee (26), as stated in the text, using MacVector v16 software (MacVector, Inc, North Carolina, USA). Proteins functionally similar to NS2 and p15 were sought using profile hidden Markov models deployed by the HMMER server (https://www.ebi.ac.uk/Tools/hmmer/). Fastq sequences in NCBI accession PRJEB28510 were downloaded to a local database and searched for sequences homologous to MKPV using MacVector v16 software.

