## Supplemental Figures for "MKPV (aka MuCPV) and related chapparvoviruses are nephro-tropic and encode novel accessory proteins p15 and NS2"

### Figure S1

(A) Overview of RACE procedure. (B) Sanger sequence traces for major RACE products.

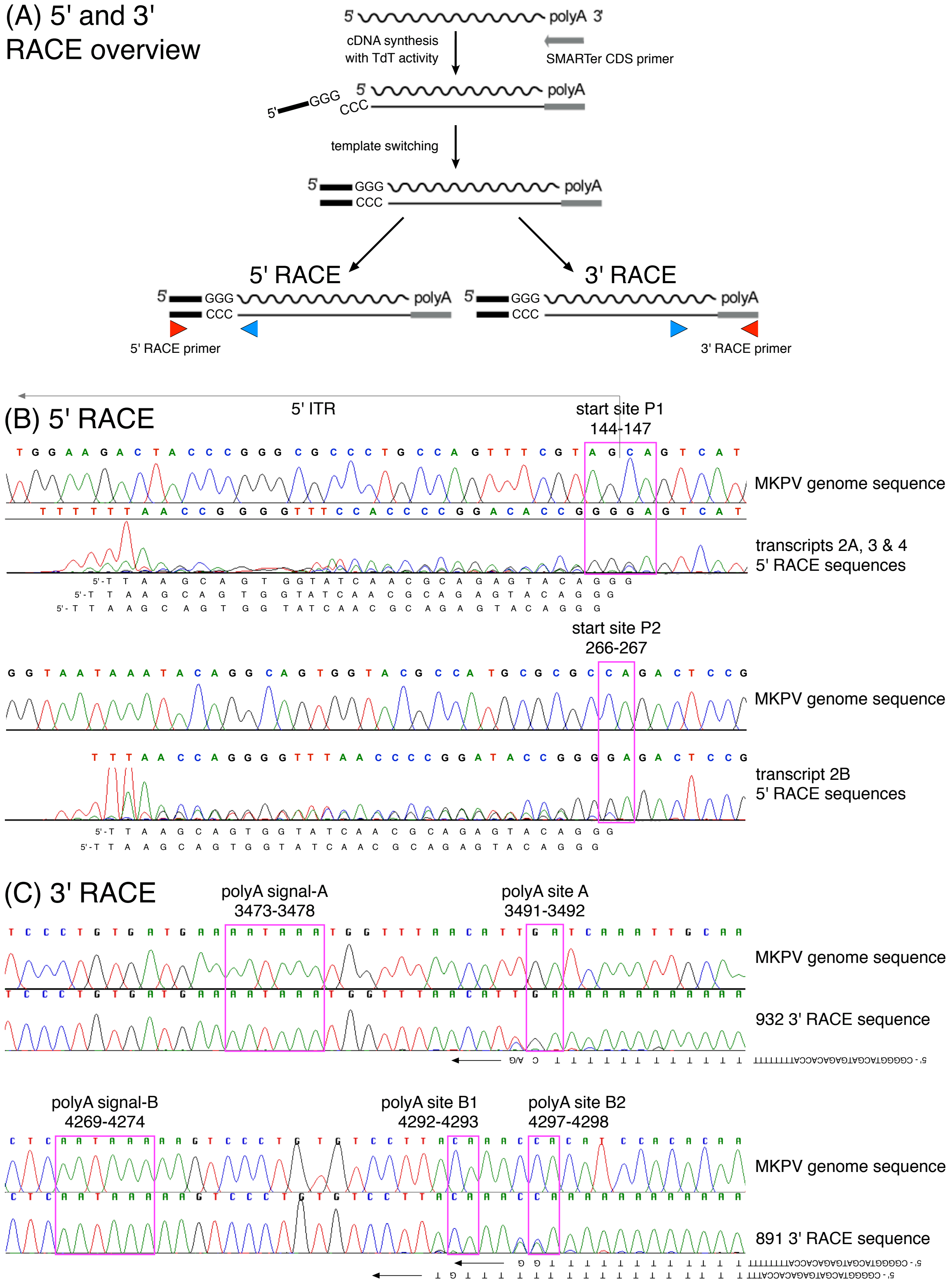

### Figure S2

Variant 3’-ITR structure created by a single-base insertion in the 3’ITR of a sub-strain of CI-MKPV (indicated by “▲” in Fig 1A).

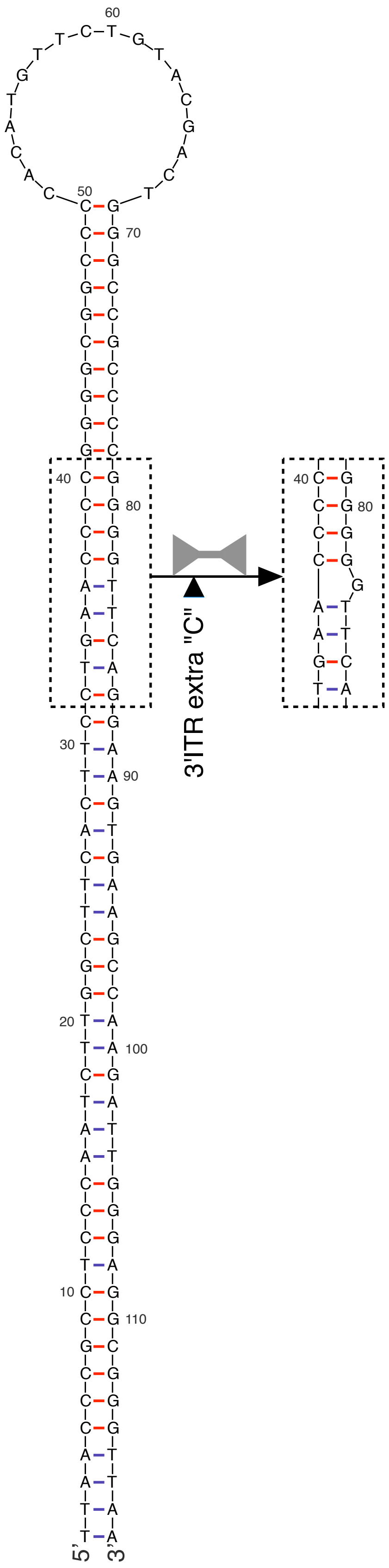

### Figure S3

Pipeline for assembly of full-length CKPV genome sequence.

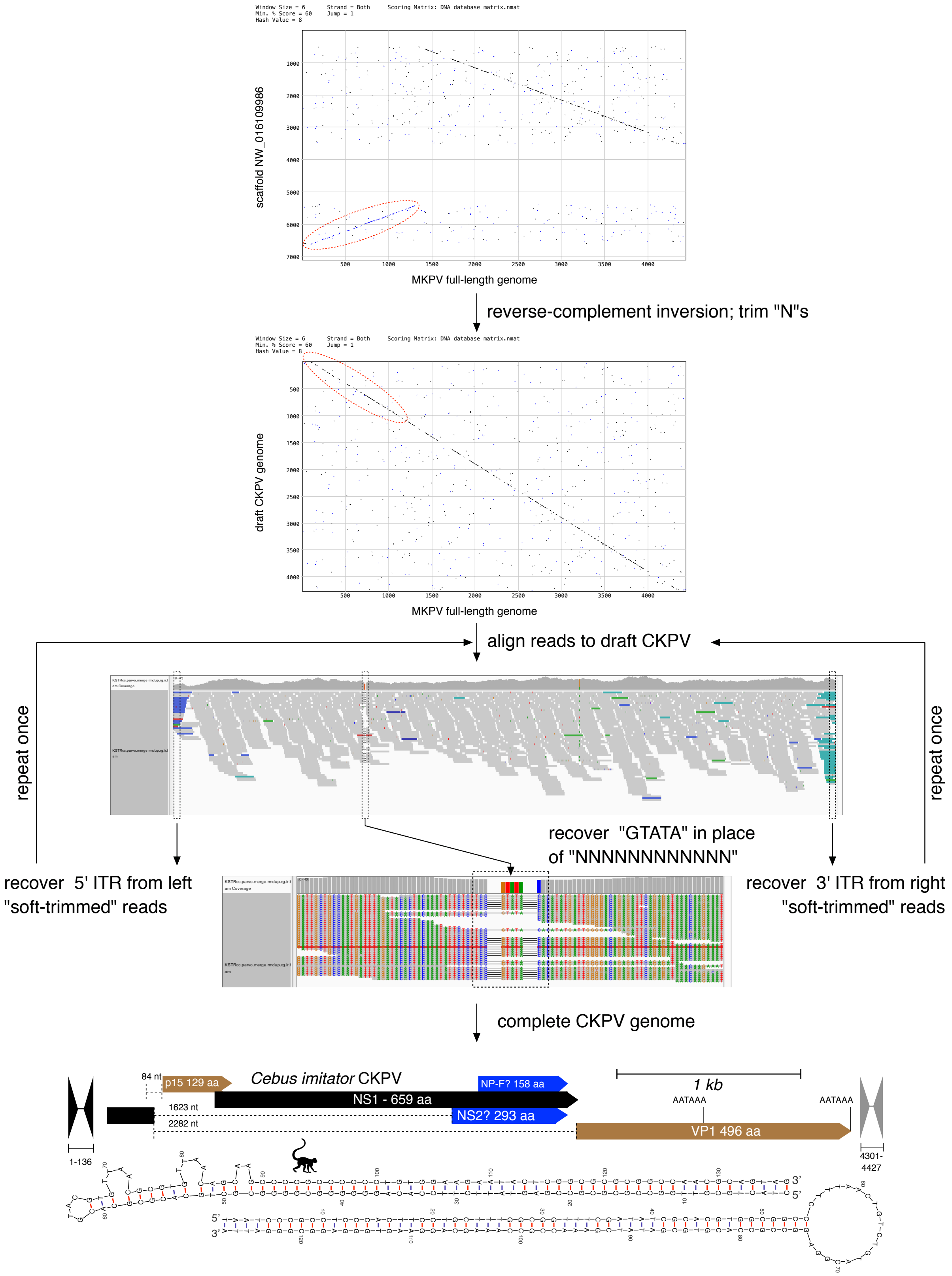

### Figure S4

Introns mapped in MKPV by RT-PCR, and corresponding hypothetical introns in related chapparvovirus species; shown red. Poly-pyrimidine tracts are underlined (with percentage pyrimidines underneath). Start ATG codons are shown in bold; STOP codons for NS1 are boxed. The splice site score generated by Genie software (ref 18) is shown above each actual (MKPV) or hypothetical (all others) splice donor and acceptor site; NS = no score generated.

**Spliced NS2**

```
Mus musculus MKPV - MH670587
Splice score:   0.99                                                            0.58
TCTCCAACTGCCGAAGGTAATTAAAAACCTTTTTTTATATCTTCTTACAGA.....ATATGCATGATACACCATTTATTTGCAGAGCTAGTGAATATAGCTGCCAAT
 S  P  T  A  E  >                                 1622 bp             67% Pyr      <E  L  V  N  I  A  A  N
```

```
Cebus imitator CKPV - KV391748
Splice score:   0.94                                                            0.95
TCACCAACACCTGAAGGTACTTATCTTTTTATTTCTTTTATGCATGCAGAT.....ATATGCACGAAACACCTTTCTTTTGCAGAACTAGTGAACATAGCTGCCAAT
 S  P  T  P  E  >                                 1629 bp             73% Pyr      <E  L  V  N  I  A  A  N
```

```
Sarcophilus harrisii CPV 2 - MK513529
Splice score:   0.99                                                            0.96
TCGCCTACATCAGAAGGTAATTATATTTACTCTTACTTTTCTTTCTAGATG.....ATATGCATGATACTCCTTTTGTTCTCAGAGCTAGTGAACATAGCTGCAAAT
 S  P  T  S  E  >                                 1623 bp             80% Pyr      <E  L  V  N  I  A  A  N
```

```
Desmodus rotundus parvovirus strain DRA25 - NC_032097
Splice score:   1.00                                                            0.51
GAACCCACAGCAGAAGGTAAGCTAACATTTGTATTATTTTTCATAGATGTC.....ACATGCATAACACCACTTTCACTTACAGAGTTAGTGAATATAGCTGCGAAT
 E  P  T  A  E  >                                 1636 bp             73% Pyr      <E  L  V  N  I  A  A  N
```

```
Ratus norvegicus parvovirus 2 isolate 9 - KX272741
Splice score:   0.89                                                            0.80
TCTCCTACGCCTGAAGGTATTTATTCTTTTTGTTTTTCTCTCTTTTAGAAT.....ATGATTGCACTGGTCTCTATTCTTGTAGAGCTAGTGAACATAGCTGCGAAT
 S  P  T  P  E  >                                 1612 bp             80% Pyr      <E  L  V  N  I  A  A  N
```

```
Eidolon helvum isolate BtPV/CMR/2014 - MG693107
Splice score:   0.89                                                            0.93
GATTCTGAACCTGAAGGTATTTATTTTTATTTTATAGATGTCTCAGTGGAC.....ACACTTCCTCTGGTATTTTGATTTATAGAAACAGCTGCACGAGCTGCAGCT
 D  S  E  P  E  >                                 1633 bp             67% Pyr      <E  T  A  A  R  A  A  A
```

```
Protobothrops mucrosquamatus ChPV NW_015402345
Splice score:   0.90                                                              NS
AGTCCATCAGCAGAAGGTAATGAAACCTTTTCTTTTAGCATTTCAGGCTAT.....ACATGATGTACCCTGAATTTACACCTAGAACAAGCGAACATAGCTGCGAAT
 S  P  S  A  E  >                                 1621 bp             60% Pyr      <E  Q  A  N  I  A  A  N
```

```
Sus scrofa PV7 isolate 42 - KU563733
Splice score:   0.98                                                            0.93
AAAGACAACTACGCAGGTGAGTGTGTAACTGCTTTTGCTTTCGCTTTAGAT.....ACTGTACTGACGGTGTGTTCGTTCGCAGATCCAGCGGTAGCTGCTGCCAAT
 K  D  N  Y  A  >                                 1663 bp             60% Pyr      <D  P  A  V  A  A  A  N
```

```
Mesitornis unicolor ChPV - NW_010204294
Splice score:   0.99                                                              NS
CCTCCTACAGCTGAGGGTAAGCAATGGCGTATTCTGGCGGTTTTTCGGTGC.....AATGCGCAAACGAACATTATACTCCTAGAGCTCGTGAACATAGCTGCAAAT
 P  P  T  A  E  >                                 1637 bp             53% Pyr      <E  L  V  N  I  A  A  N
```

```
Grus japonicus PV isolate yc-9 - KY312548
Splice score:   0.59                                                            0.50
CCAGCAACAGCTGAAGGTACAACATGTCTTTTCTGGGCTTTTCATTATTAA.....CTTCAAAAGATGCTGTATATGTTCCCAGAATTAGTGAACGAAGCTGTCAAT
 P  A  T  A  E  >                                 1640 bp             60% Pyr      <E  L  V  N  E  A  V  N
```

```
Gallus gallus PV strain RS/BR/2S - MG846442
Splice score:   1.00                                                            0.94
TCAGATCCTGAAGAAGGTACCAGTCATAATGCACAAGGTATGTCATTATTT.....ATATGTCTGATGGTATTTTTCATCCTAGAGTTACTGAATCAGGCTGCGAAT
 S  D  P  E  E  >                                 1685 bp             67% Pyr      <E  L  L  N  Q  A  A  N
```

```
Sarcophilus harrisii CPV 6 - MK513533
Splice score:   0.99                                                            0.49
CCTGAGGACCCAGCGGGTACGTATGGCAGGTCTAGGCTTTACCCTATTGCT.....AACCGAGCTGTGAGTGCGGTCCTTGCAGAGAGAGCAGAGGCAGCCAGACAA
 P  E  D  P  A  >                                 1666 bp             53% Pyr      <E  R  A  E  A  A  R  Q
```

## VP1

*Mus musculus* MKPV - MH670587

```
Splice score:   0.99                                                    0.91
TCTCCAACTGCCGAAGGTAATTAAAAACCTTTTTTTATATCTTCTTACAGA.....ATGGTTCACTTATCTATCTTATTTACAGAAACACTATGGCTGAAGATGTCA
                >                                 2289 bp          80% Pyr       <         M  A  E  D  V
```

*Cebus imitator* CKPV - KV391748

```
Splice score:   0.94                                                    0.81
TCACCAACACCTGAAGGTACTTATCTTTTTATTTCTTTTATGCATGCAGAT.....ATGGCTTGCTTATCTCTCTTATTTACAGCAACAATATGGCTGAAGATATCA
                >                                 2288 bp          87% Pyr       <         M  A  E  D  I
```

*Sarcophilus harrisii* CPV 2 - MK513529

```
Splice score:   0.99                                                    0.99
TCGCCTACATCAGAAGGTAATTATATTTACTCTTACTTTTCTTTCTAGATG.....ATGGTATTCTTATCTTTCCTATTTACAGACTCACTATGGCTGAAGATATAT
                >                                 2290 bp          80% Pyr       <         M  A  E  D  I
```

*Desmodus rotundus* parvovirus strain DRA25 - NC_032097

```
Splice score:   1.00                                                    0.89
GAACCCACAGCAGAAGGTAAGCTAACATTTGTATTATTTTTCATAGATGTC.....ATGGTATATGTATCTCTCCTACCTACAGACAAAATATGGCTGAAGATGTCA
                >                                 2330 bp          80% Pyr       <         M  A  E  D  V
```

*Ratus norvegicus* parvovirus 2 isolate 9 - KX272741

```
Splice score:   0.89                                                    0.46
TCTCCTACGCCTGAAGGTATTTATTCTTTTTGTTTTTCTCTCTTTTAGAAT.....GCTAATAAATGTAATCTTTGATTTACAGAATAAACATGGCAACTGATGTTT
                >                                 2310 bp          67% Pyr       <         M  A  T  D  V
```

*Eidolon helvum* isolate BtPV/CMR/2014 - MG693107

```
Splice score:   0.89                                                    0.95
GATTCTGAACCTGAAGGTATTTATTTTTATTTTATAGATGTCTCAGTGGAC.....CTGGTTATCTTACTTGTCTTATTTACAGACCAATTATGGCTGAAGATGTTT
                >                                 2354 bp          80% Pyr       <         M  A  E  D  V
```

*Protobothrops mucrosquamatus* ChPV NW_015402345

```
Splice score:   0.90                                                    0.85
AGTCCATCAGCAGAAGGTAATGAAACCTTTTCTTTTAGCATTTCAGGCTAT.....ATGGAAATGTTATTTGTCTTACTTGCAGAATAAATATGGCTACTGATCATG
                >                                 2285 bp          80% Pyr       <         M  A  T  D  H
```

*Sus scrofa* PV7 isolate 42 - KU563733

```
Splice score:   0.98                                0.66
AAGACAACTACGCAGGTGAGTGTGTAACTGCTTT.....CACTGGTGTGCTTATCTGTCCTATCTAGAGCATTGCTTCGGTGATCAATAAAGCGCCTGAGCTCAATCATGGCAGAACACATCACC
               >                 2343 bp          73% Pyr       <                                                M  A  E  H  I  T
```

*Mesitornis unicolor* ChPV - NW_010204294

```
Splice score:   0.99                                                NS
CCTCCTACAGCTGAGGGTAAGCAATGGCGTATTCTGGCGGTTTTTC.....GCATGTACCTCTTAAATCTGATTGGCAGCAATACCTCAGCTACATCTACCACGCATATGGCTGAAGATTATACTA
                >                          2307 bp          60% Pyr       <                                      M  A  E  D  Y  T
```

*Grus japonicus* PV isolate yc-9 - KY312548

```
Splice score: 0.59                                     0.41
CCAGCAACAGCTGAAGGTACAACATGTCTTTTCTGGGCTTTTCA.....ATTACCTATGTATGTTCCTTTAAAACAGGATTGGGAAGAATACCTGAATTTCCTGTATCATATGTATGGCTGAAGACGTCTCATA
               >                          2286 bp          60% Pyr       <                                                M  A  E  D  V  S  Y
```

*Gallus gallus* PV strain RS/BR/2S - MG846442

```
Splice score: 1.00                                                                                    0.76
TCAGATCCTGAAGAAGGTACCAGTCATAATGCACAAGGTATGTCATTATTTTCTTATATTTCAGAG.....TTTATCTGCATACGAAGATCTACATTCAGATTCAGAATGACTGAAACAATAAAAT
                >                          2398 bp                                           53% Pyr       < M  T  E  T  I  K
```

*Sarcophilus harrisii* CPV 6 - MK513533

```
Splice score:   0.99                                0.98
CCTGAGGACCCAGCGGGTACGTATGGCAGGTCTAGGC.....CATAACACTATCTATTCCTACTCCACAGGACTGGAAGAATTACTTGTCTTACTTGTATCACATTTATCACGATGCCTGAATACACTAATTGTT
               >                2186 bp          80% Pyr       <                                                      M  P  E  Y  T  N  C
```

**NS1 or p15**

*Mus musculus* MKPV - MH670587
```
Splice score:   0.99                                           0.98
CTAGAAGGAGAAGGAGGTGAGTCAGAATCGGA.....ATTAAAAACCTTTTTTATATCTTCTTACAGATGTCTATGTGGTCA
                >              88/86 bp              87% Pyr      <
                                                             p15>   M  S  M  W  S
```

*Cebus imitator* CKPV - KV391748
```
Splice score:   1.00                                           0.55
CTAGAAACCGAAAGAGGTGAGTCGCCAGACTC.....ACTTATCTTTTTATTTCTTTTATGCATGCAGATGTCTATGTGGACG
                >                84 bp               73% Pyr      <
                                                         p15> (+2) <M  S  M  W  T
```

*Sarcophilus harrisii* CPV 2 - MK513529
```
Splice score:   0.99                                           0.99
CGTATAGAGACAGACGGTGAGTCAACTCCTGA.....TAATTATATTTACTCTTACTTTTCTTTCTAGATGTCAATGTGGTCT
                >                 86 bp              93% Pyr      <
                                                             p15>   M  S  M  W  S
```

*Desmodus rotundus* parvovirus strain DRA25 - NC_032097
```
Splice score:   0.99                                           1.00
CGAACAGACGAAAAGGGTGAGTCTTCAGACGA.....GGTAAGCTAACATTTGTATTATTTTTCATAGATGTCTAGCGGCTGG
                >                84 bp               80% Pyr      <
                                                             p15>   M  S  S  G  W
```

*Ratus norvegicus* parvovirus 2 isolate 9 - KX272741
```
Splice score:   0.84                                          0.99
GGGACACAGTTTCCTGGTAGGCGTCAGTTGCG.....ATTTATTCTTTTTGTTTTTCTCTCTTTTAGAATGTCCTGGGGAAAT
                >                262 bp              93% Pyr     <
                                                        p15> (+71) <M  S  W  G  N
```

*Eidolon helvum* isolate BtPV/CMR/2014 - MG693107
```
Splice score:   0.96                                          0.99
CTGCTAAACAACCAAGGTTAGACGACGGGGAG.....GAACCTGAAGGTATTTATTTTTATTTTATAGATGTCTCAGTGGACG
                >                86 bp               80% Pyr      <
                                                             p15>   M  S  Q  W  T
```

*Protobothrops mucrosquamatus* ChPV NW_015402345
```
Splice score:   0.95                                        0.97
   <CTTTCGGAAATGGTACGCAGCGCGTGAT.....AATGAAACCTTTTCTTTTAGCATTTCAGGCTATGGCGTCTGCATGG
                >                188 bp           80% Pyr      <
                                                       p15> (+10)  <M  A  S  A  W
```

*Sus scrofa* PV7 isolate 42 - KU563733
```
Splice score:   0.67                                           1.00
GCGGACACTCGAGGAGGTAGGGGTCCGGGTCC.....GAGTGTGTAACTGCTTTTGCTTTCGCTTTAGATGGCTGGCGCGAATG
                >                145 bp               80% Pyr     <
                                                             p15>   M  A  G  A  N
```

*Mesitornis unicolor* ChPV - NW_010204294
```
Splice score:   0.95                              0.46
AGTCTGCTTCTATCAGGTAATGCCT.....CCCGCTCTCGAAGATCCTCCTACAGCTGAGGGTAAGCAATGGCGTATTCTGGCG
                >         187 bp         67% Pyr     <
                                                             p15>   M  A  Y  S  G
```

*Grus japonicus* PV isolate yc-9 - KY312548
```
Splice score:    0.99                                       0.96
TAGATTAAAAAAAGCTGGTAAGTGGCATGGAGATAAACCTCCAACTCATTCTGTTCCTCCATTAGAAGAAGGCGA.....CAACAGCTGAAGGTACAACATGTCTTTTCTGG
                 >               48 bp            80% Pyr      <
                                                                                            p15>   M  S  F  L
```

*Gallus gallus* PV strain RS/BR/2S - MG846442
```
Splice score:    0.99                                         0.96
CTAGGGTATAAGTATGAGTAAGTACA.....ATGCACAAGGTATGTCATTATTTTCTTATATTTCAGAGTATCTATGGCGGGCTTTGGAT
                 >               231 bp             80% Pyr      <
                                                                   p15>   M  A  G  F  G
```

*Sarcophilus harrisii* CPV 6 - MK513533
```
Splice score:   0.96                                   0.40
CCTGAGGACCCAGCGGGTACGAGACAGCC.....GGGACCCCAAGAGCGCTCGTCCTTAGACCCCAGAGGCCT.....CCCAGCGGGTACGTATGGCAGGTCTAGGCT
                >           253 bp          53% Pyr      <                          p15>   M  A  G  L  G
```
