## Supplemental Tables for "MKPV (aka MuCPV) and related chapparvoviruses are nephro-tropic and encode novel accessory proteins p15 and NS2"

### Table S1

PCR primers used in this study

| MKPV primers | Sequence (5'–3') | MKPV bases (MH670587) |
|---|---|---|
| 869 | GCAACACAACTGTAAATCGG | 1458–1477 |
| 870 | TGGATAATCGTGCCTAAACC | 1534–1553 |
| 889 | GGGCAGAGAGACTACGTCGGGC | 832–853 |
| 890 | CACGCAGTGCGCGCACATGC | 75–94 |
| 891 | GGTCAGATACAGAACAGGAGG | 3991–4011 |
| 893 | ACATGTTCTGTACGACTGGG | 4372–4391 |
| 900 | ATTCACTAGCTCTTCGGCAG | 478–486:2109–2119 |
| 902 | TTAGAGAGCAGGCAGATAGC | 602–621 |
| 904 | ATTGAATTCATATGCCGGGCGGAGGTA | 214–234 |
| 905 | TGTGCGGCCGCTTAAGGTGCCCAGATCCC | 2711–2728 |
| 932 | AGGCACTACCGCTTTCACTG | 3061–3080 |
| 933 | GAAAGCTTTGTAATCTCCTGTG | 3163–3184 |
| 934 | CACCTCACAGAGATCAGATGC | 1119–1139 |
| 935 | TATGCATCTGTGAGGTGGTC | 1407–1426 |
| 940 | AGCCGCATCTATGCACCAAC | 3911–3930 |
| 947 | GACGTGCCAGATCTTACTGAACC | 348–370 |
| 948 | GTGACATCTTCAGCCATAGTG | 2779–2799 |
| 955 | CTGCCAGTTTCGTAGCAGTC | 131–150 |
| **DrPV-1 primers** | | **DrPV-1 bases (NC032097)** |
| Chap-DRPv-fwd | GCAAGCGCAAGTGGAACG | 654:671 |
| Chap-DRPv-rev | GACCATCCTCCTCCTGTAG | 1418:1436 |
| **RACE primers** | | |
| SMARTer CDS primer | CGGGGTACGATGAGACACCATTTTTTTTTTTTTTTTTTTTTVN | |
| Template Switch oligo | AAGCAGTGGTATCAACGCAGAGTACAGGG | |
| 5' RACE primer | TTAAGCAGTGGTATCAACGCAGAGTACAG | |
| 3' RACE primer | CGGGGTACGATGAGACACCA | |

### {MATRIX SCIENCE} MASCOT Search Results

**Table S2**

**Re-analysis for MKPV peptides in datset PXD010540**

#### Protein View: BR004

##### sp|BR004|MKPV_p15

| | |
|---|---|
| **Database:** | BSA |
| **Score:** | 816 |
| **Nominal mass ($M_r$):** | 14673 |
| **Calculated pI:** | 7.77 |

Sequence similarity is available as an NCBI BLAST search of BR004 against nr.

##### Search parameters

| | |
|---|---|
| **MS data file:** | 6cg_Rag_01.mgf |
| **Enzyme:** | Trypsin: cuts C-term side of KR unless next residue is P. |
| **Variable modifications:** | Oxidation (M) |

##### Protein sequence coverage: 64%

Matched peptides shown in ***bold red***.

```
  1 MSMWSGPTGF SLLLISSHAN AQELLEDAIC LLSNRWHIEF EIKNQDNVWY
 51 AWGKQSRFTV GESTLQRALG DLYSQELVTF QKGPPDTSFD SALRYQKCKR
101 KWNVPDVVSL PSDDTGGAAT PVISFRKKAR
```

Unformatted sequence string: 130 residues (for pasting into other applications).

Sort peptides by ◉ Residue Number ○ Increasing Mass ○ Decreasing Mass

Show predicted peptides also

| Query | Start – End | Observed | Mr(expt) | Mr(calc) | Delta | M | Score | Expect | Rank | U | Peptide |
|---|---|---|---|---|---|---|---|---|---|---|---|
| 7350 | 36 – 43 | 367.8782 | 1100.6127 | 1100.5655 | 0.0472 | 0 | 3 | 0.5 | 1 | U | R.WHIEFEIK.N |
| 7351 | 36 – 43 | 367.8784 | 1100.6135 | 1100.5655 | 0.0481 | 0 | 3 | 0.53 | 1 | U | R.WHIEFEIK.N |
| 7352 | 36 – 43 | 551.3142 | 1100.6137 | 1100.5655 | 0.0483 | 0 | 1 | 0.82 | 1 | U | R.WHIEFEIK.N |
| 15406 | 44 – 54 | 690.8191 | 1379.6236 | 1379.6258 | -0.0022 | 0 | 27 | 0.0019 | 1 | U | K.NQDNVWYAWGK.Q |
| 15407 | 44 – 54 | 690.8193 | 1379.6240 | 1379.6258 | -0.0018 | 0 | 24 | 0.0039 | 1 | U | K.NQDNVWYAWGK.Q |
| 15408 | 44 – 54 | 690.8195 | 1379.6244 | 1379.6258 | -0.0015 | 0 | 29 | 0.0013 | 1 | U | K.NQDNVWYAWGK.Q |
| 15410 | 44 – 54 | 690.8209 | 1379.6272 | 1379.6258 | 0.0014 | 0 | 17 | 0.02 | 1 | U | K.NQDNVWYAWGK.Q |
| 15411 | 44 – 54 | 690.8213 | 1379.6280 | 1379.6258 | 0.0022 | 0 | 37 | 0.00018 | 1 | U | K.NQDNVWYAWGK.Q |
| 8398 | 58 – 67 | 568.8537 | 1135.6928 | 1136.5826 | -0.8897 | 0 | 2 | 0.59 | 1 | U | R.FTVGESTLQR.A |
| 8400 | 58 – 67 | 568.8538 | 1135.6930 | 1136.5826 | -0.8896 | 0 | 6 | 0.25 | 1 | U | R.FTVGESTLQR.A |
| 8402 | 58 – 67 | 568.8542 | 1135.6938 | 1136.5826 | -0.8887 | 0 | 5 | 0.34 | 1 | U | R.FTVGESTLQR.A |
| 8417 | 58 – 67 | 569.2978 | 1136.5810 | 1136.5826 | -0.0015 | 0 | 55 | 3.3e-06 | 1 | U | R.FTVGESTLQR.A |
| 8419 | 58 – 67 | 569.2981 | 1136.5816 | 1136.5826 | -0.0009 | 0 | 57 | 2.2e-06 | 1 | U | R.FTVGESTLQR.A |
| 8420 | 58 – 67 | 569.2983 | 1136.5821 | 1136.5826 | -0.0005 | 0 | 44 | 4.3e-05 | 1 | U | R.FTVGESTLQR.A |
| 8421 | 58 – 67 | 569.2986 | 1136.5827 | 1136.5826 | 0.0002 | 0 | 44 | 4e-05 | 1 | U | R.FTVGESTLQR.A |
| 8422 | 58 – 67 | 569.2991 | 1136.5836 | 1136.5826 | 0.0011 | 0 | 33 | 0.0005 | 1 | U | R.FTVGESTLQR.A |
| 23306 | 68 – 82 | 856.4077 | 1710.8008 | 1710.8828 | -0.0820 | 0 | 0 | 0.93 | 1 | U | R.ALGDLYSQELVTFQK.G |
| 23312 | 68 – 82 | 856.4477 | 1710.8808 | 1710.8828 | -0.0020 | 0 | 38 | 0.00017 | 1 | U | R.ALGDLYSQELVTFQK.G |
| 23313 | 68 – 82 | 856.4485 | 1710.8825 | 1710.8828 | -0.0003 | 0 | 41 | 7.8e-05 | 1 | U | R.ALGDLYSQELVTFQK.G |
| 23314 | 68 – 82 | 856.4486 | 1710.8826 | 1710.8828 | -0.0002 | 0 | 41 | 8.2e-05 | 1 | U | R.ALGDLYSQELVTFQK.G |
| 23315 | 68 – 82 | 571.3020 | 1710.8842 | 1710.8828 | 0.0013 | 0 | 22 | 0.0059 | 1 | U | R.ALGDLYSQELVTFQK.G |
| 23316 | 68 – 82 | 856.4497 | 1710.8847 | 1710.8828 | 0.0019 | 0 | 32 | 0.00057 | 1 | U | R.ALGDLYSQELVTFQK.G |
| 23317 | 68 – 82 | 856.4510 | 1710.8874 | 1710.8828 | 0.0046 | 0 | 24 | 0.0039 | 1 | U | R.ALGDLYSQELVTFQK.G |
| 23321 | 68 – 82 | 856.4667 | 1710.9188 | 1710.8828 | 0.0360 | 0 | 2 | 0.64 | 1 | U | R.ALGDLYSQELVTFQK.G |
| 35505 | 68 – 94 | 985.7258 | 2954.1556 | 2954.4662 | -0.3106 | 1 | 2 | 0.58 | 1 | U | R.ALGDLYSQELVTFQKGPPDTSFDSALR.Y |
| 35506 | 68 – 94 | 985.7263 | 2954.1570 | 2954.4662 | -0.3092 | 1 | 1 | 0.83 | 1 | U | R.ALGDLYSQELVTFQKGPPDTSFDSALR.Y |
| 12121 | 83 – 94 | 631.3284 | 1260.6422 | 1261.5939 | -0.9516 | 0 | 4 | 0.41 | 1 | U | K.GPPDTSFDSALR.Y |
| 12137 | 83 – 94 | 631.3741 | 1260.7336 | 1261.5939 | -0.8602 | 0 | 12 | 0.058 | 1 | U | K.GPPDTSFDSALR.Y |
| 12143 | 83 – 94 | 631.7101 | 1261.4057 | 1261.5939 | -0.1882 | 0 | 9 | 0.11 | 1 | U | K.GPPDTSFDSALR.Y |
| 12144 | 83 – 94 | 631.7101 | 1261.4057 | 1261.5939 | -0.1882 | 0 | 0 | 0.9 | 2 | U | K.GPPDTSFDSALR.Y |
| 12147 | 83 – 94 | 631.7103 | 1261.4060 | 1261.5939 | -0.1879 | 0 | 5 | 0.33 | 1 | U | K.GPPDTSFDSALR.Y |
| 12148 | 83 – 94 | 631.7111 | 1261.4076 | 1261.5939 | -0.1862 | 0 | 3 | 0.53 | 1 | U | K.GPPDTSFDSALR.Y |
| 12152 | 83 – 94 | 631.8026 | 1261.5906 | 1261.5939 | -0.0032 | 0 | 51 | 8.4e-06 | 1 | U | K.GPPDTSFDSALR.Y |
| 12153 | 83 – 94 | 631.8032 | 1261.5918 | 1261.5939 | -0.0021 | 0 | 44 | 3.6e-05 | 1 | U | K.GPPDTSFDSALR.Y |
| 12154 | 83 – 94 | 631.8036 | 1261.5926 | 1261.5939 | -0.0012 | 0 | 54 | 4.3e-06 | 1 | U | K.GPPDTSFDSALR.Y |
| 12155 | 83 – 94 | 631.8041 | 1261.5936 | 1261.5939 | -0.0002 | 0 | 62 | 6e-07 | 1 | U | K.GPPDTSFDSALR.Y |
| 12156 | 83 – 94 | 631.8041 | 1261.5936 | 1261.5939 | -0.0002 | 0 | 46 | 2.7e-05 | 1 | U | K.GPPDTSFDSALR.Y |
| 12157 | 83 – 94 | 631.8044 | 1261.5942 | 1261.5939 | 0.0004 | 0 | 48 | 1.7e-05 | 1 | U | K.GPPDTSFDSALR.Y |
| 12158 | 83 – 94 | 631.8047 | 1261.5948 | 1261.5939 | 0.0010 | 0 | 80 | 8.9e-09 | 1 | U | K.GPPDTSFDSALR.Y |
| 12159 | 83 – 94 | 631.8055 | 1261.5964 | 1261.5939 | 0.0026 | 0 | 81 | 8.6e-09 | 1 | U | K.GPPDTSFDSALR.Y |
| 12176 | 83 – 94 | 421.5745 | 1261.7016 | 1261.5939 | 0.1077 | 0 | 0 | 1 | 1 | U | K.GPPDTSFDSALR.Y |
| 12183 | 83 – 94 | 631.8718 | 1261.7290 | 1261.5939 | 0.1351 | 0 | 9 | 0.12 | 1 | U | K.GPPDTSFDSALR.Y |
| 12184 | 83 – 94 | 632.2184 | 1262.4222 | 1261.5939 | 0.8284 | 0 | 2 | 0.68 | 1 | U | K.GPPDTSFDSALR.Y |
| 12187 | 83 – 94 | 632.2448 | 1262.4750 | 1261.5939 | 0.8812 | 0 | 7 | 0.2 | 1 | U | K.GPPDTSFDSALR.Y |
| 12195 | 83 – 94 | 632.2925 | 1262.5704 | 1261.5939 | 0.9766 | 0 | 4 | 0.38 | 1 | U | K.GPPDTSFDSALR.Y |
| 12221 | 83 – 94 | 632.3622 | 1262.7099 | 1261.5939 | 1.1160 | 0 | 19 | 0.012 | 1 | U | K.GPPDTSFDSALR.Y |
| 22680 | 83 – 97 | 841.4058 | 1680.7970 | 1680.8107 | -0.0138 | 1 | 1 | 0.89 | 1 | U | K.GPPDTSFDSALRYQK.C |

| Query | Start – End | Observed | Mr(expt) | Mr(calc) | Delta | M | Score | Expect | Rank | U | Peptide |
|---|---|---|---|---|---|---|---|---|---|---|---|
| 22707 | 83 – 97 | 841.9218 | 1681.8290 | 1680.8107 | 1.0183 | 1 | 2 | 0.64 | 1 | U | K.GPPDTSFDSALRYQK.C |
| 33706 | 102 – 126 | 867.4013 | 2599.1821 | 2599.2919 | -0.1098 | 0 | 3 | 0.49 | 1 | U | K.WNVPDVVSLPSDDTGGAATPVISFR.K |
| 33707 | 102 – 126 | 867.4023 | 2599.1851 | 2599.2919 | -0.1068 | 0 | 7 | 0.22 | 1 | U | K.WNVPDVVSLPSDDTGGAATPVISFR.K |
| 33708 | 102 – 126 | 867.4023 | 2599.1852 | 2599.2919 | -0.1067 | 0 | 3 | 0.53 | 1 | U | K.WNVPDVVSLPSDDTGGAATPVISFR.K |
| 33711 | 102 – 126 | 867.4354 | 2599.2843 | 2599.2919 | -0.0076 | 0 | 12 | 0.068 | 1 | U | K.WNVPDVVSLPSDDTGGAATPVISFR.K |
| 33713 | 102 – 126 | 1300.6537 | 2599.2928 | 2599.2919 | 0.0010 | 0 | 0 | 0.95 | 1 | U | K.WNVPDVVSLPSDDTGGAATPVISFR.K |
| 33714 | 102 – 126 | 1300.6540 | 2599.2934 | 2599.2919 | 0.0016 | 0 | 5 | 0.32 | 1 | U | K.WNVPDVVSLPSDDTGGAATPVISFR.K |
| 33715 | 102 – 126 | 867.4386 | 2599.2940 | 2599.2919 | 0.0021 | 0 | 50 | 1e-05 | 1 | U | K.WNVPDVVSLPSDDTGGAATPVISFR.K |
| 33717 | 102 – 126 | 1300.6547 | 2599.2948 | 2599.2919 | 0.0030 | 0 | 1 | 0.74 | 1 | U | K.WNVPDVVSLPSDDTGGAATPVISFR.K |
| 33718 | 102 – 126 | 867.4391 | 2599.2955 | 2599.2919 | 0.0036 | 0 | 30 | 0.00092 | 1 | U | K.WNVPDVVSLPSDDTGGAATPVISFR.K |
| 33719 | 102 – 126 | 867.4392 | 2599.2956 | 2599.2919 | 0.0038 | 0 | 28 | 0.0016 | 1 | U | K.WNVPDVVSLPSDDTGGAATPVISFR.K |
| 33720 | 102 – 126 | 867.4394 | 2599.2965 | 2599.2919 | 0.0047 | 0 | 22 | 0.0065 | 1 | U | K.WNVPDVVSLPSDDTGGAATPVISFR.K |

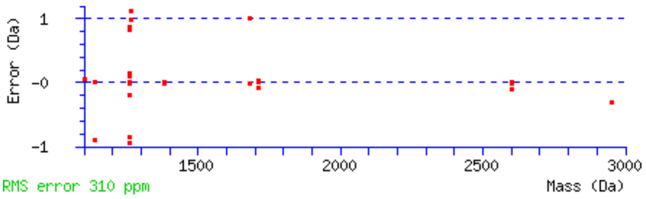

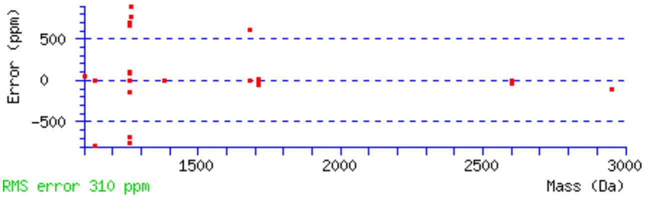

Mascot: http://www.matrixscience.com/

### MASCOT Search Results

#### Protein View: BR002

##### sp|BR002|MKPV_NS1

| | |
|---|---|
| **Database:** | BSA |
| **Score:** | 219 |
| **Nominal mass ($M_r$):** | 73424 |
| **Calculated pI:** | 7.06 |

Sequence similarity is available as **an NCBI BLAST search of BR002 against nr**.

##### Search parameters

| | |
|---|---|
| **MS data file:** | 6cg_Rag_01.mgf |
| **Enzyme:** | Trypsin: cuts C-term side of KR unless next residue is P. |
| **Variable modifications:** | **Oxidation (M)** |

##### Protein sequence coverage: 29%

Matched peptides shown in ***bold red***.

```
  1 MQAQMERARR SLSALRRYWW GGNACHQLSE ESEIISPENL KQIMLNWDSR
 51 VWQACVLGIW DTVPVRDPRP YCFLLTNIPS VKKWLICAEE DSNEQTHIHL
101 LALTSQRSDA FKRTLEKTWK QVAIVAMSDI EEPDPTLEIV KCQKCHKPSS
151 LLAYMAKDPH WIAANDMQTL GIFESVYAHD WGQRFREKQT LDKAKKTDPT
201 TSQMHTITAE ITEVIMQHNC KSVEDCMKAA PTVIAKHLHR AGLGTIIQNC
251 ISWVTATGGG WSLPSIGAKH PPEPEAIHTI LLHQGISPAD FDPIFYKWLA
301 KEETKKNTLV LWGPSNTGKS AFISGLKTCT NWGEVVNSNT FAFEALINAQ
351 LGVWEEPLIS PELAEKAKQI FEGMETSIPV KYRKPVKLPR IPIIITTNHA
401 PWRFCTKEEE MFRNRMYIFT WSQNMHDTPF ICRASEYSCQ CRVCQTSRGG
451 QACAGGQSAG SLQRKEQSVS ELVQPEPSSS YVSTRSLPVS REETPLPAAE
501 GLGSHHQRHC SSPGGESIER THSPRPSCST GSSTSDSLRP SGEHRSSDPG
551 AGISCSFSGS LECVESPLSG GDDGDDLPRD RMGEPTSPDS STGSSDISRP
601 RGKRRHSQEM VVLGETQSKK TRDQVSTAVT GMGRDLGTLN IPTRAQWFTY
651 LSYLQKHYG
```

Unformatted sequence string: **659 residues** (for pasting into other applications).

Sort peptides by ◉ Residue Number ○ Increasing Mass ○ Decreasing Mass

Show predicted peptides also

| Query | Start – End | Observed | Mr(expt) | Mr(calc) | ppm | M | Score | Expect | Rank | U | Peptide |
|---|---|---|---|---|---|---|---|---|---|---|---|
| 7855 | 1 – 9 | 560.7870 | 1119.5594 | 1119.5277 | 28.4 | 1 | 1 | 0.74 | 1 | U | -.MQAQMERAR.R |
| 8381 | 1 – 9 | 568.7719 | 1135.5292 | 1135.5226 | 5.84 | 1 | 1 | 0.78 | 1 | U | -.MQAQMERAR.R + Oxidation (M) |
| 8382 | 1 – 9 | 568.7731 | 1135.5317 | 1135.5226 | 8.01 | 1 | 1 | 0.83 | 1 | U | -.MQAQMERAR.R + Oxidation (M) |
| 8385 | 1 – 9 | 379.5247 | 1135.5523 | 1135.5226 | 26.1 | 1 | 3 | 0.52 | 1 | U | -.MQAQMERAR.R + Oxidation (M) |
| 4922 | 2 – 9 | 503.2555 | 1004.4964 | 1004.4821 | 14.2 | 1 | 3 | 0.48 | 1 | U | M.QAQMERAR.R + Oxidation (M) |
| 9060 | 2 – 10 | 581.2877 | 1160.5608 | 1160.5832 | -19.4 | 2 | 0 | 0.95 | 1 | U | M.QAQMERARR.S + Oxidation (M) |
| 5399 | 8 – 16 | 515.3058 | 1028.5970 | 1028.6203 | -22.6 | 2 | 4 | 0.38 | 1 | U | R.ARRSLSALR.R |
| 5401 | 8 – 16 | 515.3064 | 1028.5982 | 1028.6203 | -21.4 | 2 | 2 | 0.62 | 1 | U | R.ARRSLSALR.R |
| 36202 | 194 – 221 | 1052.8326 | 3155.4760 | 3155.5413 | -20.7 | 2 | 0 | 0.9 | 1 | U | K.AKKTDPTTSQMHTITAEITEVIMQHNCK.S |
| 36294 | 194 – 221 | 1063.5410 | 3187.6012 | 3187.5312 | 22.0 | 2 | 2 | 0.65 | 1 | U | K.AKKTDPTTSQMHTITAEITEVIMQHNCK.S + 2 Oxidation (M) |
| 36295 | 194 – 221 | 1063.5410 | 3187.6012 | 3187.5312 | 22.0 | 2 | 0 | 0.9 | 1 | U | K.AKKTDPTTSQMHTITAEITEVIMQHNCK.S + 2 Oxidation (M) |
| 36296 | 194 – 221 | 1063.5435 | 3187.6087 | 3187.5312 | 24.3 | 2 | 6 | 0.23 | 1 | U | K.AKKTDPTTSQMHTITAEITEVIMQHNCK.S + 2 Oxidation (M) |
| 36297 | 194 – 221 | 797.9099 | 3187.6105 | 3187.5312 | 24.9 | 2 | 3 | 0.49 | 1 | U | K.AKKTDPTTSQMHTITAEITEVIMQHNCK.S + 2 Oxidation (M) |
| 36298 | 194 – 221 | 1063.5464 | 3187.6174 | 3187.5312 | 27.0 | 2 | 0 | 0.97 | 1 | U | K.AKKTDPTTSQMHTITAEITEVIMQHNCK.S + 2 Oxidation (M) |
| 37452 | 196 – 228 | 1261.2407 | 3780.7003 | 3780.7137 | -3.55 | 2 | 0 | 0.92 | 1 | U | K.KTDPTTSQMHTITAEITEVIMQHNCKSVEDCMK.A + 2 Oxidation (M) |
| 20323 | 222 – 236 | 789.8872 | 1577.7598 | 1577.7793 | -12.3 | 1 | 10 | 0.11 | 1 | U | K.SVEDCMKAAPTVIAK.H + Oxidation (M) |
| 20331 | 222 – 236 | 789.9122 | 1577.8099 | 1577.7793 | 19.4 | 1 | 2 | 0.6 | 1 | U | K.SVEDCMKAAPTVIAK.H + Oxidation (M) |
| 804 | 229 – 236 | 385.7422 | 769.4698 | 769.4698 | 0.099 | 0 | 17 | 0.021 | 1 | U | K.AAPTVIAK.H |
| 4910 | 298 – 305 | 502.7794 | 1003.5443 | 1003.5338 | 10.5 | 1 | 1 | 0.77 | 1 | U | K.WLAKEETK.K |
| 15611 | 307 – 319 | 693.8731 | 1385.7315 | 1385.7303 | 0.89 | 0 | 28 | 0.0014 | 1 | U | K.NTLVLWGPSNTGK.S |
| 22628 | 367 – 381 | 839.4602 | 1676.9058 | 1676.8807 | 15.0 | 1 | 1 | 0.82 | 1 | U | K.AKQIFEGMETSIPVK.Y |
| 18349 | 369 – 381 | 747.9013 | 1493.7880 | 1493.7436 | 29.8 | 0 | 0 | 0.96 | 1 | U | K.QIFEGMETSIPVK.Y + Oxidation (M) |
| 7604 | 408 – 415 | 555.7415 | 1109.4684 | 1109.4924 | -21.5 | 1 | 4 | 0.42 | 1 | U | K.EEEMFRNR.M |
| 7605 | 408 – 415 | 555.7420 | 1109.4694 | 1109.4924 | -20.6 | 1 | 5 | 0.3 | 1 | U | K.EEEMFRNR.M |
| 31928 | 465 – 485 | 779.7243 | 2336.1511 | 2336.1496 | 0.64 | 1 | 40 | 0.00011 | 1 | U | R.KEQSVSELVQPEPSSSYVSTR.S |
| 25557 | 492 – 508 | 457.9796 | 1827.8895 | 1827.8864 | 1.70 | 0 | 48 | 1.5e-05 | 1 | U | R.EETPLPAAEGLGSHHQR.H |
| 26486 | 604 – 619 | 629.0100 | 1884.0080 | 1883.9636 | 23.6 | 2 | 1 | 0.87 | 1 | U | K.RRHSQEMVVLGETQSK.K |
| 26489 | 604 – 619 | 472.0100 | 1884.0109 | 1883.9636 | 25.1 | 2 | 3 | 0.5 | 1 | U | K.RRHSQEMVVLGETQSK.K |
| 26493 | 604 – 619 | 472.0104 | 1884.0123 | 1883.9636 | 25.9 | 2 | 4 | 0.44 | 1 | U | K.RRHSQEMVVLGETQSK.K |
| 26494 | 604 – 619 | 629.0118 | 1884.0136 | 1883.9636 | 26.5 | 2 | 8 | 0.17 | 1 | U | K.RRHSQEMVVLGETQSK.K |
| 20572 | 606 – 619 | 794.8740 | 1587.7334 | 1587.7563 | -14.4 | 0 | 3 | 0.46 | 1 | U | R.HSQEMVVLGETQSK.K + Oxidation (M) |
| 20573 | 606 – 619 | 794.8755 | 1587.7364 | 1587.7563 | -12.5 | 0 | 0 | 0.93 | 1 | U | R.HSQEMVVLGETQSK.K + Oxidation (M) |
| 27809 | 606 – 622 | 987.4865 | 1972.9584 | 1973.0000 | -21.1 | 2 | 3 | 0.55 | 1 | U | R.HSQEMVVLGETQSKKTR.D + Oxidation (M) |
| 17964 | 621 – 634 | 739.8859 | 1477.7572 | 1477.7307 | 17.9 | 1 | 1 | 0.81 | 2 | U | K.TRDQVSTAVTGMGR.D |
| 18338 | 621 – 634 | 747.8624 | 1493.7102 | 1493.7257 | -10.4 | 1 | 6 | 0.27 | 1 | U | K.TRDQVSTAVTGMGR.D + Oxidation (M) |

| Query | Start – End | Observed | Mr(expt) | Mr(calc) | ppm | M | Score | Expect | Rank | U | Peptide |
|---|---|---|---|---|---|---|---|---|---|---|---|
| 10928 | 623 – 634 | 611.2980 | 1220.5815 | 1220.5820 | -0.37 | 0 | 53 | 5.2e-06 | 1 | U | R.DQVSTAVTGMGR.D |
| 11386 | 623 – 634 | 619.2938 | 1236.5730 | 1236.5769 | -3.09 | 0 | 5 | 0.34 | 1 | U | R.DQVSTAVTGMGR.D + Oxidation (M) |
| 31762 | 623 – 644 | 773.3859 | 2317.1358 | 2317.1696 | -14.6 | 1 | 1 | 0.75 | 1 | U | R.DQVSTAVTGMGRDLGTLNIPTR.A + Oxidation (M) |
| 7284 | 635 – 644 | 550.3072 | 1098.5998 | 1098.6033 | -3.16 | 0 | 35 | 0.0003 | 1 | U | R.DLGTLNIPTR.A |
| 7285 | 635 – 644 | 550.3076 | 1098.6006 | 1098.6033 | -2.43 | 0 | 48 | 1.5e-05 | 1 | U | R.DLGTLNIPTR.A |
| 7287 | 635 – 644 | 550.3083 | 1098.6020 | 1098.6033 | -1.16 | 0 | 41 | 8.7e-05 | 1 | U | R.DLGTLNIPTR.A |
| 35595 | 635 – 659 | 995.8366 | 2984.4880 | 2984.5185 | -10.2 | 2 | 0 | 0.97 | 1 | U | R.DLGTLNIPTRAQWFTYLSYLQKHYG.- |
| 19589 | 645 – 656 | 774.3951 | 1546.7756 | 1546.7820 | -4.11 | 0 | 0 | 0.91 | 1 | U | R.AQWFTYLSYLQK.H |
| 26822 | 645 – 659 | 952.9606 | 1903.9066 | 1903.9257 | -10.0 | 1 | 0 | 0.97 | 1 | U | R.AQWFTYLSYLQKHYG.- |
| 26824 | 645 – 659 | 952.9645 | 1903.9144 | 1903.9257 | -5.91 | 1 | 1 | 0.83 | 1 | U | R.AQWFTYLSYLQKHYG.- |

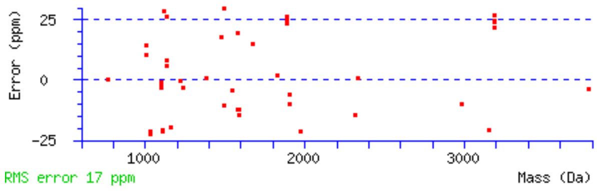

Mascot: http://www.matrixscience.com/

### MASCOT Search Results

#### Protein View: BR003

##### sp|BR003|MKPV_VP1

**Database:** BSA
**Score:** 26
**Nominal mass ($M_r$):** 57134
**Calculated pI:** 6.32

Sequence similarity is available as an NCBI BLAST search of BR003 against nr.

##### Search parameters

**MS data file:** 6cg_Rag_01.mgf
**Enzyme:** Trypsin: cuts C-term side of KR unless next residue is P.
**Variable modifications:** Oxidation (M)

##### Protein sequence coverage: 25%

Matched peptides shown in ***bold red***.

```
  1 MAEDVTFHNT YMVYWKNQPF IYPNTNINPP NAHTMSAGAI NTGWHIIPTI
 51 LWKHFLTPKQ WTEFTINYEA YTVKGYSCTI YNPIPMTQQL AIQGTTAFTA
101 FNNTIYTLGA QDDLYETAYH NWYSDDSTGD YKAFNLSFKE GQYKNLSGSW
151 KKTIWPIYSW RTENARNASS STYSYLNGID SYAVWPRTKD KELIPTGVFW
201 DPLNDANGIL ELRPGKNSMS FSWEQHPCDE NKWFNIDQIA KWFPYTVDTP
251 YLNPQTYGPP GSYKLYGEDD PDQLTTPSSW TAYSAKNDYT IPNLLDMPIV
301 PMQWFWQEIQ KSIAEVPDVK KPMLYWAGTE YECYKYGPTQ CFLKGIPLFD
351 DNDTHVATTT QGCFRISLHL AGKKRRSRIY APTWGPLSWR QCYATDTPFA
401 PSMVRYRTGG ARRTWTNINR DAEGVHKDFH YREDPYDITS TVPDTRGTAT
451 VTDSKATMHP YEQAASGMYL NHKEMRQVRA AAEATRSQPA VAMQTQ
```

Unformatted sequence string: 496 residues (for pasting into other applications).

Sort peptides by ◉ Residue Number ○ Increasing Mass ○ Decreasing Mass

Show predicted peptides also

| Query | Start – End | Observed | Mr(expt) | Mr(calc) | ppm | M | Score | Expect | Rank | U | Peptide |
|---|---|---|---|---|---|---|---|---|---|---|---|
| 1587 | 133 – 139 | 413.7190 | 825.4235 | 825.4385 | -18.2 | 0 | 4 | 0.41 | 1 | | K.AFNLSFK.E |
| 1588 | 133 – 139 | 413.7195 | 825.4244 | 825.4385 | -17.0 | 0 | 3 | 0.48 | 1 | | K.AFNLSFK.E |
| 3208 | 145 – 152 | 460.2473 | 918.4801 | 918.4923 | -13.2 | 1 | 1 | 0.87 | 1 | | K.NLSGSWKK.T |
| 3212 | 145 – 152 | 460.2492 | 918.4838 | 918.4923 | -9.20 | 1 | 4 | 0.4 | 1 | | K.NLSGSWKK.T |
| 26697 | 321 – 335 | 949.4015 | 1896.7884 | 1896.8426 | -28.6 | 0 | 1 | 0.78 | 1 | | K.KPMLYWAGTEYECYK.Y + Oxidation (M) |
| 6053 | 336 – 344 | 528.7692 | 1055.5238 | 1055.5110 | 12.1 | 0 | 0 | 0.9 | 1 | | K.YGPTQCFLK.G |
| 6054 | 336 – 344 | 528.7693 | 1055.5240 | 1055.5110 | 12.4 | 0 | 0 | 0.91 | 1 | | K.YGPTQCFLK.G |
| 31682 | 345 – 365 | 770.0301 | 2307.0685 | 2307.0590 | 4.09 | 0 | 1 | 0.74 | 1 | | K.GIPLFDDNDTHVATTTQGCFR.I |
| 7970 | 366 – 375 | 374.9007 | 1121.6804 | 1121.7033 | -20.4 | 2 | 2 | 0.64 | 1 | | R.ISLHLAGKKR.R |
| 17155 | 379 – 390 | 723.8697 | 1445.7249 | 1445.7456 | -14.3 | 0 | 1 | 0.84 | 1 | | R.IYAPTWGPLSWR.Q |
| 6152 | 413 – 420 | 530.7847 | 1059.5548 | 1059.5574 | -2.43 | 1 | 3 | 0.47 | 1 | | R.RTWTNINR.D |
| 6154 | 413 – 420 | 530.7852 | 1059.5558 | 1059.5574 | -1.50 | 1 | 4 | 0.42 | 1 | | R.RTWTNINR.D |
| 28930 | 456 – 473 | 683.6467 | 2047.9182 | 2047.9244 | -3.03 | 0 | 25 | 0.0032 | 1 | U | K.ATMHPYEQAASGMYLNHK.E |
| 28932 | 456 – 473 | 683.6492 | 2047.9258 | 2047.9244 | 0.68 | 0 | 14 | 0.038 | 1 | U | K.ATMHPYEQAASGMYLNHK.E |
| 29116 | 456 – 473 | 688.9799 | 2063.9179 | 2063.9193 | -0.69 | 0 | 13 | 0.051 | 1 | U | K.ATMHPYEQAASGMYLNHK.E + Oxidation (M) |
| 6483 | 477 – 486 | 536.8040 | 1071.5934 | 1071.5785 | 13.9 | 1 | 8 | 0.15 | 1 | U | R.QVRAAAEATR.S |
| 29754 | 477 – 496 | 1057.5599 | 2113.1052 | 2113.0698 | 16.8 | 2 | 1 | 0.86 | 1 | U | R.QVRAAAEATRSQPAVAMQTQ.- |
| 29755 | 477 – 496 | 1057.5605 | 2113.1064 | 2113.0698 | 17.3 | 2 | 2 | 0.58 | 1 | U | R.QVRAAAEATRSQPAVAMQTQ.- |
| 23676 | 480 – 496 | 577.6382 | 1729.8929 | 1729.8417 | 29.6 | 1 | 1 | 0.75 | 1 | U | R.AAAEATRSQPAVAMQTQ.- |

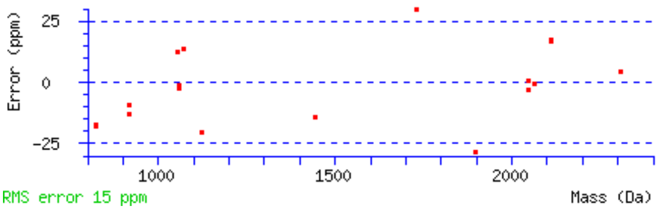

**Mascot: http://www.matrixscience.com/**

### MASCOT Search Results

#### Protein View: BR008

##### sp|BR008|MKPV_p10|89aa (151-420)

**Database:** BSA
**Score:** 227
**Nominal mass ($M_r$):** 9815
**Calculated pI:** 9.68

Sequence similarity is available as an NCBI BLAST search of BR008 against nr.

##### Search parameters

**MS data file:** 6cg_Rag_01.mgf
**Enzyme:** Trypsin: cuts C-term side of KR unless next residue is P.
**Variable modifications:** Oxidation (M)

##### Protein sequence coverage: 66%

Matched peptides shown in ***bold red***.

```
 1 MCCCPLCGCD  CGDTEQNILI  RLICRAEVIN  TGSGTPCAPD  SAQQASGTRI
51 GRVCRDCLKQ  LEEHKKTCQI  LLNLIRLTLQ  IHQLQSDRV
```

Unformatted sequence string: 89 residues (for pasting into other applications).

Sort peptides by ◉ Residue Number ○ Increasing Mass ○ Decreasing Mass

Show predicted peptides also

| Query | Start | – End | Observed | Mr(expt) | Mr(calc) | ppm | M | Score | Expect | Rank | U | Peptide |
|---|---|---|---|---|---|---|---|---|---|---|---|---|
| 34817 | 19 | – 49 | 935.1343 | 2802.3811 | 2802.3389 | 15.1 | 1 | 3 | 0.45 | 1 | U | R.LICRAEVINTGSGTPCAPDSAQQASGTR.I |
| 34818 | 19 | – 49 | 935.1362 | 2802.3867 | 2802.3389 | 17.1 | 1 | 2 | 0.57 | 1 | U | R.LICRAEVINTGSGTPCAPDSAQQASGTR.I |
| 1739 | 53 | – 59 | 418.7211 | 835.4277 | 835.4044 | 27.9 | 1 | 1 | 0.71 | 1 | U | R.VCRDCLK.Q |
| 9825 | 67 | – 76 | 593.8429 | 1185.6712 | 1185.6903 | -16.1 | 0 | 2 | 0.66 | 1 | U | K.TCQILLNLIR.L |
| 19649 | 77 | – 89 | 775.9360 | 1549.8574 | 1549.8576 | -0.12 | 1 | 58 | 1.7e-06 | 1 | U | R.LTLQIHQLQSDRV.- |
| 19650 | 77 | – 89 | 775.9360 | 1549.8575 | 1549.8576 | -0.069 | 1 | 42 | 6.8e-05 | 1 | U | R.LTLQIHQLQSDRV.- |
| 19651 | 77 | – 89 | 517.6267 | 1549.8583 | 1549.8576 | 0.41 | 1 | 81 | 7.4e-09 | 1 | U | R.LTLQIHQLQSDRV.- |
| 19652 | 77 | – 89 | 517.6271 | 1549.8595 | 1549.8576 | 1.19 | 1 | 85 | 2.8e-09 | 1 | U | R.LTLQIHQLQSDRV.- |

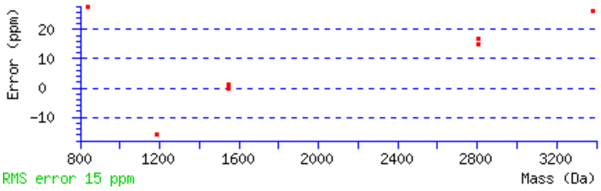

Mascot: http://www.matrixscience.com/

### Table S3

Summary of significant MKPV peptides in independent LC-MS/MS dataset PXDnnnn

| Protein Group | Accession | -10lgP | Coverage (%) | Area Sample 1 | #Peptides | #Unique | #Spec Sample 1 | PTM | Avg. Mass |
|---|---|---|---|---|---|---|---|---|---|
| 203 | MKPV_p15 | 145.69 | 63 | 1.50E+09 | 12 | 12 | 34 | | 14683 |
| 362 | MKPV_NS1 | 152.56 | 28 | 1.91E+08 | 14 | 14 | 22 | | 73471 |
| 1584 | MKPV_VP1 | 53.89 | 10 | 1.59E+07 | 4 | 4 | 4 | Deamidation (NQ) | 57170 |
| 1770 | MKPV_p10 | 35.76 | 16 | 2.20E+07 | 2 | 2 | 3 | | 9815 |

| Protein Group | Protein Accession | Peptide | Unique | -10lgP | Mass | Length | ppm | m/z | z | RT | Area Sample 1 | Fraction | Scan | Source File | #Feature | #Feature Sample 1 | Start | End | PTM | AScore |
|---|---|---|---|---|---|---|---|---|---|---|---|---|---|---|---|---|---|---|---|---|
| 203 | MKPV_p15 | K.GPPDTSFDSALR.Y | Y | 57.38 | 1261.5939 | 12 | -2.5 | 631.8026 | 2 | 55.06 | 4.86E+08 | 1 | 20336 | Infected_01.raw | 1 | 1 | 83 | 94 | | |
| 203 | MKPV_p15 | R.ALGDLYSQELVTFQK.G | Y | 53.78 | 1710.8828 | 15 | 1.6 | 856.4501 | 2 | 80.66 | 2.28E+08 | 1 | 33880 | Infected_01.raw | 3 | 3 | 68 | 82 | | |
| 203 | MKPV_p15 | K.NQDNVWYAWGK.Q | Y | 48.61 | 1379.6259 | 11 | 0.4 | 690.8205 | 2 | 72.58 | 3.76E+07 | 1 | 29917 | Infected_01.raw | 1 | 1 | 44 | 54 | | |
| 203 | MKPV_p15 | R.KWNVPDVVSLPSDDTGGAATPVISFR.K | Y | 42.49 | 2727.3867 | 26 | 0.5 | 910.1366 | 3 | 86.37 | 6.13E+07 | 1 | 36772 | Infected_01.raw | 1 | 1 | 101 | 126 | | |
| 203 | MKPV_p15 | R.FTVGESTLQR.A | Y | 41.61 | 1136.5825 | 10 | -0.8 | 569.2981 | 2 | 49.57 | 5.95E+08 | 1 | 17244 | Infected_01.raw | 1 | 1 | 58 | 67 | | |
| 203 | MKPV_p15 | S.DDTGGAATPVISFR.K | Y | 41.3 | 1405.6837 | 14 | -1.6 | 703.848 | 2 | 64.49 | 7.43E+06 | 1 | 25529 | Infected_01.raw | 1 | 1 | 113 | 126 | | |
| 203 | MKPV_p15 | R.WHIEFEIK.N | Y | 38.56 | 1100.5654 | 8 | 1.4 | 367.8629 | 3 | 74.99 | 2.47E+07 | 1 | 31328 | Infected_01.raw | 1 | 1 | 36 | 43 | | |
| 203 | MKPV_p15 | S.LPSDDTGGAATPVISFR.K | Y | 35.6 | 1702.8525 | 17 | -0.9 | 852.4327 | 2 | 67.62 | 7.23E+06 | 1 | 27168 | Infected_01.raw | 1 | 1 | 110 | 126 | | |
| 203 | MKPV_p15 | K.WNVPDVVSLPSDDTGGAATPVISFR.K | Y | 35.54 | 2599.2917 | 25 | -1 | 867.437 | 3 | 94.02 | 5.11E+07 | 1 | 40317 | Infected_01.raw | 2 | 2 | 102 | 126 | | |
| 203 | MKPV_p15 | N.VPDVVSLPSDDTGGAATPVISFR.K | Y | 21.74 | 2299.1694 | 23 | -0.6 | 1150.5913 | 2 | 82.81 | 2.41E+06 | 1 | 35047 | Infected_01.raw | 1 | 1 | 104 | 126 | | |
| 203 | MKPV_p15 | R.ALGDLYSQELVTF.Q | Y | 20.05 | 1454.7292 | 13 | -1.1 | 728.3711 | 2 | 96.71 | 1.87E+06 | 1 | 41486 | Infected_01.raw | 1 | 1 | 68 | 80 | | |
| 203 | MKPV_p15 | R.KWNVPDVVSLPSDDTGGAATPVISF.R | Y | 17.64 | 2571.2856 | 25 | 3.4 | 1286.6544 | 2 | 96.06 | 9.41E+05 | 1 | 41124 | Infected_01.raw | 1 | 1 | 101 | 125 | | |
| 362 | MKPV_NS1 | R.EETPLPAAEGLGSHHQR.H | Y | 57.59 | 1827.8864 | 17 | -1.5 | 457.9782 | 4 | 41.7 | 3.21E+07 | 1 | 13023 | Infected_01.raw | 2 | 2 | 492 | 508 | | |
| 362 | MKPV_NS1 | K.NTLVLWGPSNTGK.S | Y | 50.48 | 1385.7303 | 13 | -3.9 | 693.8698 | 2 | 68.87 | 1.10E+07 | 1 | 27859 | Infected_01.raw | 1 | 1 | 307 | 319 | | |
| 362 | MKPV_NS1 | K.QIFEGMETSIPVK.Y | Y | 49.41 | 1477.7487 | 13 | -1.3 | 739.8806 | 2 | 71.09 | 1.24E+07 | 1 | 29127 | Infected_01.raw | 1 | 1 | 369 | 381 | | |
| 362 | MKPV_NS1 | K.HPPEPEAIHTILLHQGISPADFDPIFYK.W | Y | 46.66 | 3181.6235 | 28 | -0.8 | 796.4125 | 4 | 87.47 | 3.36E+07 | 1 | 37445 | Infected_01.raw | 1 | 1 | 270 | 297 | | |
| 362 | MKPV_NS1 | K.DPHWIAANDMQTLGIFESVYAHDWGQR.F | Y | 45.71 | 3156.4512 | 27 | 0.2 | 790.1202 | 4 | 100.93 | 4.17E+06 | 1 | 43216 | Infected_01.raw | 2 | 2 | 158 | 184 | | |
| 362 | MKPV_NS1 | R.DQVSTAVTGMGR.D | Y | 44.84 | 1220.5819 | 12 | 0.6 | 611.2986 | 2 | 46.18 | 2.45E+07 | 1 | 15480 | Infected_01.raw | 1 | 1 | 623 | 634 | | |
| 362 | MKPV_NS1 | R.IPIIITTNHAPWR.F | Y | 44.49 | 1530.8671 | 13 | 2.3 | 511.2975 | 3 | 69.4 | 1.18E+07 | 1 | 28127 | Infected_01.raw | 2 | 2 | 391 | 403 | | |
| 362 | MKPV_NS1 | R.DLGTLNIPTR.A | Y | 31.61 | 1098.6033 | 10 | -0.6 | 550.3086 | 2 | 62.65 | 2.52E+07 | 1 | 24473 | Infected_01.raw | 1 | 1 | 635 | 644 | | |
| 362 | MKPV_NS1 | K.SAFISGLK.T | Y | 30.68 | 821.4647 | 8 | -0.8 | 411.7393 | 2 | 54.48 | 1.43E+07 | 1 | 20107 | Infected_01.raw | 1 | 1 | 320 | 327 | | |
| 362 | MKPV_NS1 | R.IPIIITTNH.A | Y | 30.37 | 1020.5968 | 9 | -0.3 | 511.3055 | 2 | 61.05 | 4.36E+06 | 1 | 23647 | Infected_01.raw | 1 | 1 | 391 | 399 | | |
| 362 | MKPV_NS1 | H.QGISPADFDPIFYK.W | Y | 29.57 | 1596.7823 | 14 | 0.9 | 799.3992 | 2 | 88.01 | 3.13E+06 | 1 | 37637 | Infected_01.raw | 1 | 1 | 284 | 297 | | |
| 362 | MKPV_NS1 | K.QIMLNWDSR.V | Y | 25.28 | 1161.5601 | 9 | -1.5 | 581.7864 | 2 | 67.44 | 5.21E+06 | 1 | 27089 | Infected_01.raw | 1 | 1 | 42 | 50 | | |
| 362 | MKPV_NS1 | R.KEQSVSELVQPEPSSSY.V | Y | 23.98 | 1892.9003 | 17 | -3.2 | 947.4543 | 2 | 57.69 | 7.75E+06 | 1 | 21777 | Infected_01.raw | 1 | 1 | 465 | 481 | | |
| 362 | MKPV_NS1 | R.MGEPTSPDSSTGSSDISRPR.G | Y | 22.04 | 2062.9226 | 20 | -2.1 | 688.6467 | 3 | 36.37 | 1.09E+06 | 1 | 10183 | Infected_01.raw | 1 | 1 | 582 | 601 | | |
| 1584 | MKPV_VP1 | K.SIAEVPDVK.K | Y | 35.45 | 956.5178 | 9 | -0.9 | 479.2657 | 2 | 47.33 | 7.63E+06 | 1 | 16005 | Infected_01.raw | 1 | 1 | 312 | 320 | | |
| 1584 | MKPV_VP1 | K.NDYTIPNLLDMPIVPMQWFWQEIQK.S | Y | 22.49 | 3118.5295 | 25 | -1.2 | 1040.5159 | 3 | 134.06 | 3.74E+05 | 1 | 56548 | Infected_01.raw | 1 | 1 | 287 | 311 | | |
| 1584 | MKPV_VP1 | K.ATMHPYEQAASGMYLN(+.98)HK.E | Y | 21.55 | 2048.9084 | 18 | -2.8 | 683.9749 | 3 | 50.6 | 4.34E+06 | 1 | 17899 | Infected_01.raw | 1 | 1 | 456 | 473 | Deamidation (NQ) | N16:Deamidation (NQ):8.51 |
| 1584 | MKPV_VP1 | K.ATMHPYEQAASGMY.L | Y | 15.06 | 1555.6436 | 14 | -2.9 | 778.8268 | 2 | 53.71 | 3.53E+06 | 1 | 19606 | Infected_01.raw | 1 | 1 | 456 | 469 | | |
| 1770 | MKPV_p10 | R.LTLQIHQLQSDR.V | Y | 24.43 | 1450.7892 | 12 | -0.4 | 484.6035 | 3 | 55.37 | 6.09E+06 | 1 | 20525 | Infected_01.raw | 1 | 1 | 97 | 108 | | |
| 1770 | MKPV_p10 | R.LTLQIHQLQSDRV | Y | 22.66 | 1549.8577 | 13 | -0.3 | 517.6263 | 3 | 61.41 | 1.59E+07 | 1 | 23827 | Infected_01.raw | 2 | 2 | 97 | 109 | | |

### Table S4

Summary of ISH in multiple tissues.

| **Animal description (1)** | | | | | | |
|---|---|---|---|---|---|---|
| **MSKCC-WCM Accession #** | 16-1653-1 | | | 15-3577-1 | | |
| **Sex** | Female | | | Female | | |
| **Age** | Unknown (adult) | | | 12 months | | |
| **Strain** | NSG | | | NSG | | |
| **Cause of death** | Inclusion body nephropathy caused by MKPV | | | Inclusion body nephropathy caused by MKPV | | |
| **Previously reported MKPV testing (1)** | | | | | | |
| **MKPV PCR on FFPE kidney tissue** | Positive | | | Positive | | |
| **MKPV ISH on FFPE kidney tissue** | Strongly positive, tubular cells, multifocal | | | NP | | |
| **ISH results** | **MKPV ISH** | **Mouse Ppib ISH (2)** | **Dapb ISH (3)** | **MKPV ISH** | **Mouse Ppib ISH** | **Dapb ISH** |
| **Heart** | Negative | NP | NP | Negative | Positive | NP |
| **Lungs** | Negative | NP | NP | Negative | Positive | NP |
| **Thymus** | U | NP | NP | Negative | Positive | NP |
| **Kidneys** | U (4) | NP | NP | Strongly positive, tubular cells, multifocal | NP | NP |
| **Liver** | Negative | NP | NP | Negative | NP | NP |
| **Gallbladder** | Negative | NP | NP | Negative | NP | NP |
| **Stomach** | Negative | NP | NP | Negative | NP | NP |
| **Duodenum** | Negative | NP | NP | Negative | NP | NP |
| **Jejunum** | Negative | NP | NP | Negative | NP | NP |
| **Ileum** | Negative | NP | NP | Negative | NP | NP |
| **Cecum** | Mildly positive, mucosal epithelium and lamina propria, multifocal | NP | NP | Negative | NP | NP |
| **Colon** | Negative | NP | NP | U | NP | NP |
| **Salivary glands** | Negative | NP | NP | Negative | NP | NP |
| **Uterus** | Negative | Positive | NP | Negative | NP | NP |
| **Urinary bladder** | Mildly positive, urothelium, multifocal; Strongly positive, casts in lumen | Positive | NP | Mildly positive, urothelium, multifocal; Strongly positive, casts in lumen | NP | NP |
| **Pancreas** | Negative | NP | NP | Negative | NP | NP |
| **Adrenals** | Negative | Positive | Negative | U | NP | NP |
| **Ovaries** | Negative | Positive | Negative | Negative | NP | NP |
| **Oviducts** | Negative | Positive | Negative | Negative | NP | NP |
| **Trachea** | Negative | Positive | Negative | Negative | Positive | NP |
| **Esophagus** | Negative | Positive | Negative | Negative | Positive | NP |
| **Thyroid** | Negative | Positive | Negative | Negative | NP | NP |
| **Skin (trunk)** | Negative | Positive | Negative | U | NP | NP |
| **Skeletal muscles** | Negative | NP | NP | Negative | NP | NP |
| **Bones (femur, tibia, sternum, vertebrae, skull)** | Not interpreted (5) | NP | NP | Not interpreted (5) | NP | NP |
| **Bone marrow (femur, tibia, sternum, vertebrae)** | Not interpreted (5) | NP | NP | Not interpreted (5) | NP | NP |
| **Stifle joint** | Not interpreted (5) | NP | NP | Not interpreted (5) | NP | NP |
| **Nerves (hind limb, spine)** | Not interpreted (5) | NP | NP | Not interpreted (5) | NP | NP |
| **Spinal cord** | Not interpreted (5) | NP | NP | Not interpreted (5) | NP | NP |
| **Oral mucosa** | Not interpreted (5) | NP | NP | Not interpreted (5) | Negative (5) | Negative |
| **Teeth** | Not interpreted (5) | NP | NP | Not interpreted (5) | Negative (5) | Negative |
| **Nasal mucosa** | Not interpreted (5) | NP | NP | Not interpreted (5) | Negative (5) | Negative |
| **Eyes** | Not interpreted (5) | NP | NP | Not interpreted (5) | Negative (5) | Negative |
| **Harderian gland** | Not interpreted (5) | NP | NP | Not interpreted (5) | Negative (5) | Negative |
| **Pituitary** | Not interpreted (5) | Negative (5) | Negative | Not interpreted (5) | NP | NP |
| **Brain** | Not interpreted (5) | Negative (5) | Negative | Not interpreted (5) | Negative (5) | Negative |
| **Ears** | Not interpreted (5) | Negative (5) | Negative | Not interpreted (5) | NP | NP |

**Notes:**

1. As described in reference 9.
2. Positive control probe.
3. Negative control probe.
4. Tissue exhausted by previous ISH staining; kidney sample from this mouse was previously found to be ISH positive as described in reference 9.
5. Due to inadequate RNA preservation caused by formic acid decalcification, as shown by negative Ppib results on all decalcified tissues tested.

NSG, NOD.Cg-*Prkdc*$^{scid}$ *Il2rg*$^{tm1Wjl}$/SzJ; FFPE, formalin-fixed paraffin-embedded; NP, not performed; U, sample unavailable.

### Table S5

IBN-postive, MKPV PCR-negative kidneys (ref 9) re-screened by ISH.

| Site[1] | Animal ID[1] | Histological evidence of IBN [1, 2] | MKPV 25-cycle PCR status[1,3] | MKPV ISH status [3] |
|---|---|---|---|---|
| MSKCC-WCM[4] | 07-2391-1 | Yes | Negative | Positive |
| MSKCC-WCM | 08-0763-1 | Yes | Negative | Positive |
| MSKCC-WCM | 08-0763-4 | Yes | Negative | Negative |
| MSKCC-WCM | 08-2405-1 | Yes | Negative | Positive |
| MSKCC-WCM | 09-2711-1 | Yes | Negative | Positive |

**Notes**
1. As previously reported (9)
2. Inclusion body nephropathy
3. Performed on formalin-fixed paraffin-embedded kidney tissue
4. Memorial Sloan Kettering Cancer Center and Weill Cornell Medicine
